# Label-free volumetric dynamic atlas of fresh mouse organs using dynamic full-field optical coherence tomography

**DOI:** 10.64898/2026.10.28.762177

**Authors:** Qiang Zhang, Mingyuan Liu, Hongyuan Cui, Shulang Lin, Chengming Wang, Jie Xie, Lihui Zhang, Hanlu Ma, Cheng Ma, Ping Xue, Bin He

## Abstract

Dynamic full-field optical coherence tomography (D-FFOCT) enables label-free visualization of tissue microarchitecture and endogenous intracellular dynamics. Here, we establish a volumetric atlas of ten fresh mouse organs using a mobile D-FFOCT imaging platform. The system enables large-area imaging of approximately 1 cm^2^ within 15 min with 1,024 frames per field of view, achieving 0.75-µm lateral resolution and imaging depths of up to 200 µm. Large-area mosaics, three-dimensional reconstructions, and comparisons with conventional histology reveal diverse organ-specific tissue architectures. Beyond morphological characterization, quantitative analysis of dynamic intensity and mean fluctuation frequency across nine organs reveals distinct dynamic signatures and tissue heterogeneity. Longitudinal imaging of brain, liver, and intestinal tissues further demonstrates organ-dependent temporal changes in endogenous dynamics following excision. Finally, we publicly release raw D-FFOCT imaging datasets from all ten organs, providing an open resource for quantitative tissue characterization, algorithm development, and reproducible research. Together, this study provides a comprehensive, openly accessible optical atlas of fresh mouse organs, establishing a foundation for systematic tissue characterization and advancing the broader application of D-FFOCT in biomedical research.

## 1. Introduction

Understanding the structural organization and functional heterogeneity of biological organs is fundamental to investigating tissue physiology, disease progression, and responses to biological interventions. Over the past decade, large-scale single-cell transcriptomic studies have established comprehensive molecular atlases across mouse organs [1], while spatial transcriptomics has further connected cellular identities with their spatial organization [2,3]. In parallel, advanced three-dimensional (3D) imaging techniques, including tissue clearing combined with light-sheet microscopy, have enabled detailed visualization of tissue architectures across multiple spatial scales [4–6]. However, these approaches generally involve tissue dissociation, sectioning, fixation, labeling, or extensive sample preparation, which may alter native tissue organization and preclude the observation of endogenous cellular dynamics. Moreover, destructive sample preparation limits the ability to acquire complementary structural, dynamic, and molecular information from the same specimen. A label-free imaging approach capable of preserving fresh tissue integrity while providing comprehensive morphological and dynamic information would therefore represent a valuable complement to existing organ atlases.

An important requirement for constructing such an atlas is the ability to visualize tissue architecture over large spatial scales without sacrificing microscopic resolution. Biological organs exhibit pronounced spatial heterogeneity, with distinct anatomical compartments, cellular populations, and functional units distributed across extended tissue regions. High-resolution microscopy can reveal fine cellular details, but its limited field of view often restricts observations to isolated regions of interest, making it difficult to interpret local structures within their broader anatomical context. Conversely, large-area imaging provides the spatial continuity necessary to identify tissue boundaries, anatomical landmarks, regional organization, and spatially distributed microstructures. The importance of preserving spatial context has been demonstrated by large-scale anatomical atlases and whole-organ imaging approaches, which enable the localization and interpretation of microscopic structures within extended tissue architectures [7,8]. This capability is particularly relevant to mouse organs, where characteristic tissue structures extend over millimeter scales, while individual cells and subcellular features require micrometer- or submicrometer-scale resolution. Combining large-area coverage with high spatial resolution therefore enables hierarchical tissue characterization, linking organ-level architecture to local cellular morphology and reducing the sampling bias associated with isolated microscopic observations. When further integrated with depth-resolved imaging, this approach can reveal the 3D organization of tissue microstructures while preserving their spatial relationships within the larger specimen.

Full-field optical coherence tomography (FFOCT) offers a promising approach by combining low-coherence interferometry with parallel camera-based detection to provide high-resolution, depth-resolved images of biological tissues without physical sectioning or exogenous labeling [9–13]. Unlike conventional histology, FFOCT enables optical sectioning directly within intact tissue specimens, allowing volumetric visualization while preserving the samples for subsequent examination. Dynamic full-field optical coherence tomography (D-FFOCT) further extends this capability by exploiting temporal fluctuations in interferometric signals associated with intracellular and subcellular motion [14]. These fluctuations provide endogenous dynamic contrast that can reveal cellular structures that are otherwise difficult to distinguish using static scattering information alone. Moreover, spectral analysis of the temporal interferometric signals enables quantitative characterization of tissue dynamics through parameters such as fluctuation intensity and characteristic frequency [14,16]. Previous studies have demonstrated D-FFOCT imaging in freshly excised tissues from several organs [14], three-dimensional live imaging of retinal organoids [15], and longitudinal monitoring of cellular organization and activity [17]. Recent developments have further expanded its applications to rapid optical histology, cancer tissue assessment, and high-resolution imaging of fresh biological tissues [18–22]. Together, these advances establish D-FFOCT as a powerful approach for combining label-free morphological visualization with endogenous dynamic information.

Despite these developments, several challenges remain in extending D-FFOCT toward systematic, multi-organ tissue characterization. Mouse organs differ substantially in cellular composition, extracellular matrix organization, optical scattering properties, and intracellular dynamics, giving rise to diverse morphological appearances and dynamic signatures. Although previous studies have demonstrated D-FFOCT in several tissue types, comprehensive and consistently acquired datasets integrating large-area structural information, depth-resolved microarchitecture, and quantitative dynamic measurements across diverse normal organs remain limited. In particular, most existing investigations have focused on tissue-specific morphological features, methodological improvements, or dynamic contrast in selected biological samples, whereas systematic quantitative comparisons of intrinsic dynamic characteristics across multiple normal organs remain relatively unexplored [14,17–22]. Dynamic signal intensity and characteristic fluctuation frequency provide complementary descriptions of tissue motion, reflecting the magnitude and temporal distribution of interferometric fluctuations, respectively. Their joint analysis may therefore reveal tissue-dependent dynamic characteristics that cannot be adequately captured by either parameter alone. Furthermore, differences in imaging configurations, acquisition conditions, and signal-processing methods can influence quantitative dynamic measurements and complicate comparisons between studies [23,24]. Establishing a consistent multi-organ imaging framework, together with accessible raw interferometric datasets, would facilitate systematic evaluation of tissue-dependent dynamic contrast and support the development of reproducible D-FFOCT analysis methods.

Beyond spatial characterization, repeated measurements of endogenous dynamic contrast offer opportunities to investigate temporal changes in fresh tissues following excision. Conventional histology primarily captures tissue morphology at a fixed endpoint, whereas D-FFOCT enables longitudinal observation of the same tissue region without exogenous labeling or destructive sample preparation. Following excision, tissues undergo progressive physiological and biochemical alterations associated with the interruption of blood supply and changes in their extracellular environment. These processes may influence intracellular motion before pronounced morphological deterioration becomes apparent. Previous investigations have demonstrated the feasibility of longitudinal D-FFOCT imaging in living cell cultures and retinal organoids [17], while other implementations of dynamic OCT have explored label-free assessment of excised organs under preservation conditions, suggesting potential applications in organ transplantation and tissue quality evaluation [25]. Nevertheless, different organs may exhibit distinct temporal responses owing to variations in cellular composition, structural organization, and physiological requirements. Characterizing these organ-dependent dynamic trajectories may therefore provide additional information for interpreting post-excision tissue changes. Equally important, the availability of raw time-resolved interferometric datasets would enable independent evaluation of dynamic parameters, imaging algorithms, and analysis strategies, facilitating further methodological development within the D-FFOCT community.

Here, we establish a label-free volumetric atlas of fresh mouse organs using an integrated, mobile D-FFOCT imaging platform. The system combines automated large-area scanning, high-resolution optical sectioning, and quantitative dynamic imaging, enabling the acquisition of approximately 1 cm^2^ of tissue within 15 min using 1,024 temporal frames per field of view, with a lateral resolution of 0.75 µm and imaging depths of up to 200 µm. The acquisition protocol also allows shorter imaging times by reducing the number of temporal frames. We systematically investigate ten freshly excised mouse organs spanning six major organ systems, including the brain, eye, thymus, inguinal lymph node, liver, pancreas, duodenum, kidney, ovary, and testis. Large-area mosaics reveal organ-specific anatomical organization and spatial heterogeneity, while local comparisons with hematoxylin and eosin (H&E) histology and 3D reconstructions demonstrate the ability to resolve tissue microarchitecture across multiple spatial scales. Beyond morphological characterization, we jointly quantify mean dynamic intensity and mean fluctuation frequency across 120 imaging locations from nine organs, revealing distinct organ-dependent dynamic signatures. We further investigate the temporal evolution of endogenous dynamics in freshly excised brain, liver, and duodenal tissues over observation periods of up to 30, 20, and 15 h, respectively. Finally, we publicly release raw D-FFOCT interferometric imaging datasets from all ten organs, providing an open resource for quantitative tissue characterization, algorithm development, and reproducible analysis. Together, this work establishes a comprehensive, openly accessible optical atlas of fresh mouse organs that integrates large-area morphology, volumetric microarchitecture, and endogenous dynamics, providing a foundation for systematic tissue characterization and broader applications of D-FFOCT in biomedical research.

## 2. Results

### 2.1 Integrated mobile D-FFOCT platform for large-area volumetric imaging

Mouse organs are relatively small, and tissue collection is often irreversible, making it desirable to obtain complementary morphological and molecular information from the same specimen. Conventional histological and molecular analysis workflows frequently require tissue sectioning, staining, or dissociation, which consume valuable samples and may introduce spatial correspondence errors when different measurements are performed on adjacent sections. To address these limitations, we developed an integrated, mobile dynamic full-field optical coherence tomography (D-FFOCT) imaging platform for rapid, label-free examination of freshly excised mouse organs (Fig. 1). The system is housed in a compact, cart-based configuration measuring approximately 55 × 67 × 100 cm³, allowing flexible deployment in animal facilities, pathology laboratories, and biomedical imaging facilities. Fresh tissue specimens can be imaged with minimal preparation, without fixation, histological sectioning, staining, or tissue clearing. This non-destructive imaging approach preserves tissue structural integrity while capturing endogenous dynamic contrast associated with intracellular and subcellular motion. The platform supports both automated large-area imaging and depth-resolved volumetric acquisition of selected tissue regions.

The imaging system is based on D-FFOCT, which combines low-coherence interferometry with temporal analysis of interferometric intensity fluctuations to visualize tissue microarchitecture and intracellular dynamics. A broadband light-emitting diode (LED) centered at 660 nm provides low-coherence illumination to a Linnik-type interferometric configuration comprising two matched water-immersion objectives. The resulting full-field interference signals are recorded using a customized high-speed (450Hz) complementary metal–oxide–semiconductor (CMOS) camera. Tissue specimens are mounted on a motorized three-axis translation stage, enabling automated lateral scanning for large-area image mosaicking and axial scanning for volumetric reconstruction. Image acquisition, stage motion, reconstruction, and dynamic signal analysis are coordinated through an integrated computing system utilizing central processing unit (CPU) and graphics processing unit (GPU) resources. Lossless data compression is incorporated to facilitate storage of large-scale time-resolved datasets, particularly during multi-organ and longitudinal imaging experiments.

The integrated platform achieves a lateral resolution of approximately 0.75 µm and enables large-area imaging of approximately 1 cm^2^ of tissue within 15 min when acquiring 1,024 temporal frames per field of view. The acquisition time can be further reduced by using fewer temporal frames, providing flexibility in balancing imaging throughput and dynamic signal characterization. Depth-resolved image stacks can be acquired over an axial range of up to 200 µm, although the effective imaging depth depends on tissue optical properties. The combination of large-area coverage and high spatial resolution allows anatomical structures spanning millimeter-scale tissue regions to be examined alongside local cellular microarchitecture. For relatively small mouse organs, the imaging area can cover a substantial portion, or even the entirety, of the exposed tissue cross-section, providing spatial context that is difficult to obtain from isolated microscopic fields of view. As illustrated in Fig. 1, representative D-FFOCT images of ten freshly excised mouse organs, including the eyeball, brain, thymus, liver, kidney, duodenum, pancreas, inguinal lymph node, testis, and ovary, reveal diverse organ-specific structural features and endogenous dynamic contrast. Importantly, the non-destructive nature of the imaging workflow allows the specimens to be retained for subsequent histological or molecular analyses, offering the possibility of correlating label-free optical information with complementary measurements from the same tissue.

**Fig. 1.**
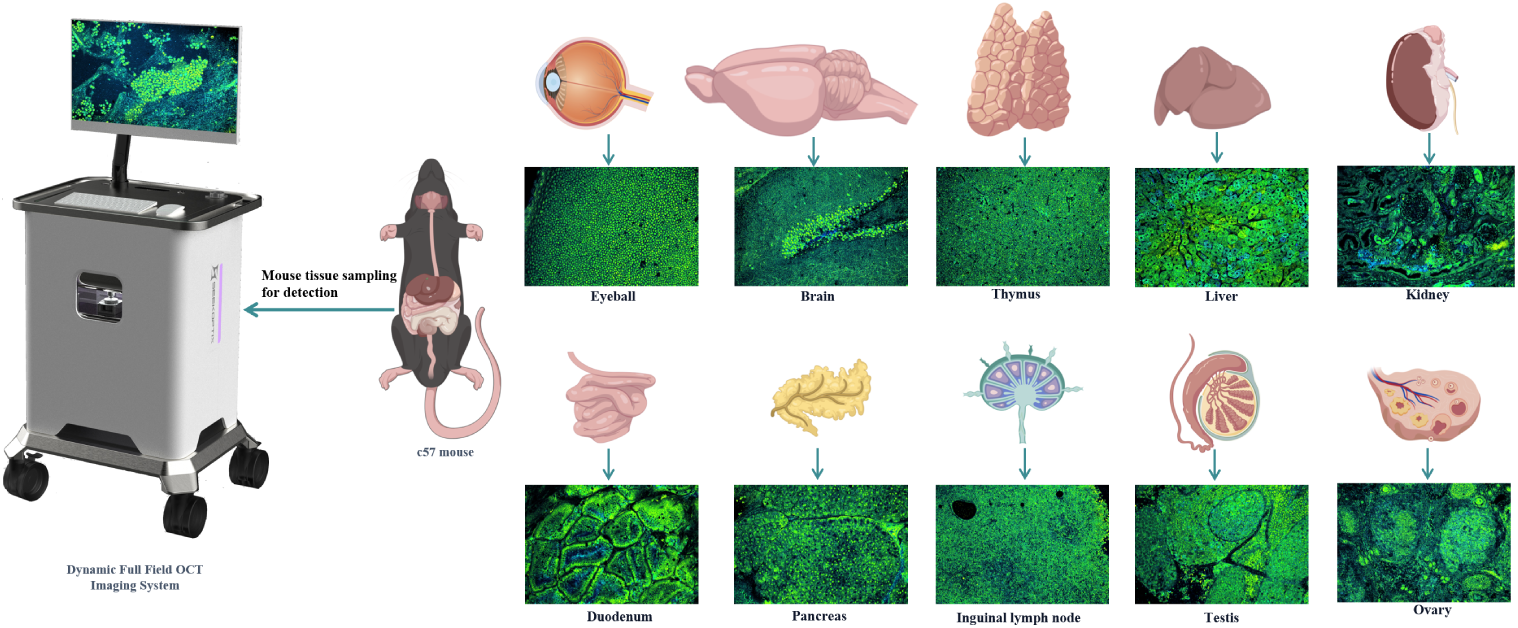
Overview of the mobile D-FFOCT imaging platform for label-free volumetric imaging of fresh mouse organs. (Left) Schematic rendering of the integrated cart-based imaging system, incorporating a dynamic full-field optical coherence tomography (D-FFOCT) module illuminated by a 660 nm LED. A motorized three-axis translation stage enables automated large-area mosaicking and depth-resolved volumetric acquisition, while an integrated computing unit performs image acquisition, reconstruction, and denoising. (Middle) Schematic illustration of tissue sampling from C57BL/6J mice. Freshly excised organs can be imaged with minimal sample preparation, without fixation, physical sectioning, staining, or tissue clearing, thereby preserving native tissue architecture and endogenous dynamic contrast. (Right) Representative label-free D-FFOCT images of ten freshly excised mouse organs, illustrating the applicability of the platform across diverse tissue types. The upper row shows the eyeball, brain, thymus, liver, and kidney, while the lower row shows the duodenum, pancreas, inguinal lymph node, testis, and ovary. The images reveal organ-specific microarchitectural features through endogenous dynamic contrast.

### 2.2 Large-area and volumetric imaging of fresh mouse organs

To evaluate the applicability of the D-FFOCT platform across diverse tissue architectures, we performed systematic imaging of ten freshly excised mouse organs without fixation, dehydration, tissue clearing, or staining (Figs. 2–11). Photographs of the fresh specimens were acquired before imaging, and corresponding tissue samples were subsequently processed for hematoxylin and eosin (H&E) staining to provide histological references for structural interpretation. Here, we extend D-FFOCT to systematic multi-organ imaging under a consistent acquisition framework, combining large-area morphological visualization with depth-resolved characterization of local tissue microarchitecture. The following sections present the imaging results for each organ, with en face images acquired in the x–y plane at selected depths beneath the tissue surface.

#### 2.2.1. Brain

The mouse brain exhibits complex anatomical organization, including distinct gray-matter regions, white-matter pathways, and highly ordered neuronal layers. Figure 2(a) presents a large-area D-FFOCT mosaic of a freshly excised mouse brain in the coronal plane, showing an overall anatomical organization consistent with the corresponding specimen photograph (Fig. 2(b)). The large-area imaging results, further illustrated in Visualization 1, reveal the spatial relationships among major anatomical structures. White-matter pathways, including the corpus callosum, internal capsule, and hippocampal fiber tracts, exhibit elongated or band-like contrast patterns, while the ventricular cavities appear as well-defined low-signal regions.

**Fig. 2.**
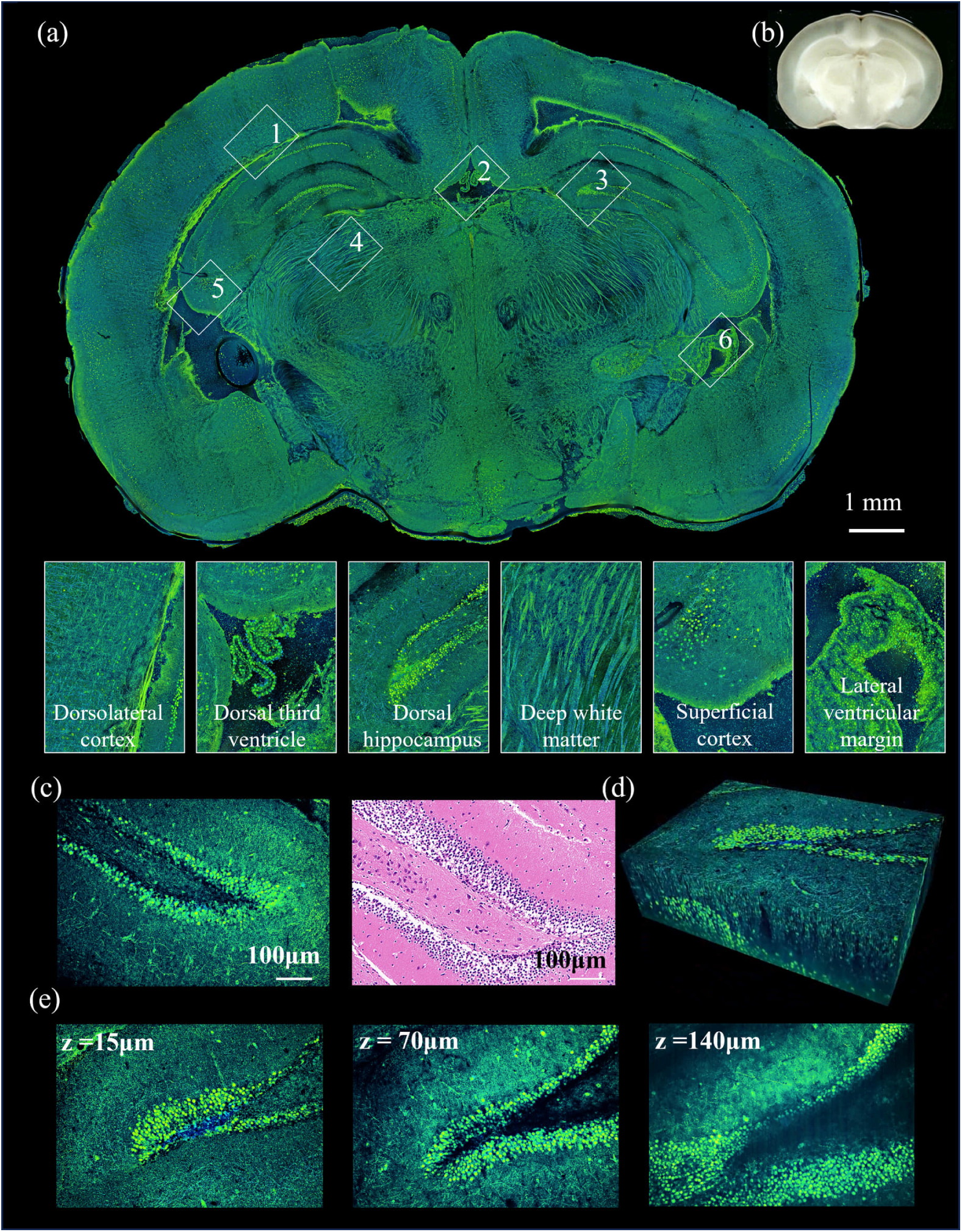
Large-area and volumetric D-FFOCT imaging of freshly excised mouse brain tissue. (a) Large-area D-FFOCT mosaic of a coronal mouse brain section, showing the overall anatomical organization and major structural features. Six numbered regions indicate representative areas for detailed examination, with region 3 corresponding to the dorsal hippocampus. (b) Photograph of the corresponding fresh brain section, showing anatomical features consistent with the D-FFOCT mosaic. (c) Magnified D-FFOCT image of region 3 and a corresponding H&E-stained histological section, highlighting the characteristic layered organization of the hippocampus, including the dentate granule cell layer. (d) Three-dimensional reconstruction of the dorsal hippocampal region. (e) Representative en face images acquired at depths of 15, 70, and 140 µm, demonstrating the continuity of hippocampal layered structures across different imaging depths. Scale bars: 1 mm (a); 100 µm (c, e).

Six representative regions were selected to illustrate regional structural diversity. These include vessel-like structures in the cerebral cortex (region 1), branching choroid plexus-like structures near the third ventricle (region 2), the characteristic layered architecture of the dorsal hippocampus (region 3), organized fiber-like structures within deep white matter (region 4), layered cortical features (region 5), and ventricular boundaries with adjacent branching structures (region 6). In particular, the dorsal hippocampus reveals elongated bands consistent with the CA pyramidal cell layers and the curved granule cell layer of the dentate gyrus.

The magnified D-FFOCT image of region 3 (Fig. 2(c)) shows layered structures that correspond morphologically to those identified in the adjacent H&E-stained section. Three-dimensional reconstruction (Fig. 2(d) and Visualization 2) and en face images acquired at depths of 15, 70, and 140 µm (Fig. 2(e)) further reveal the spatial continuity of the hippocampal layers across depth, despite variations in image contrast. These results demonstrate the ability of D-FFOCT to connect large-area brain anatomy with local depth-resolved microarchitecture.

#### 2.2.2. Eyeball

**Fig. 3.**
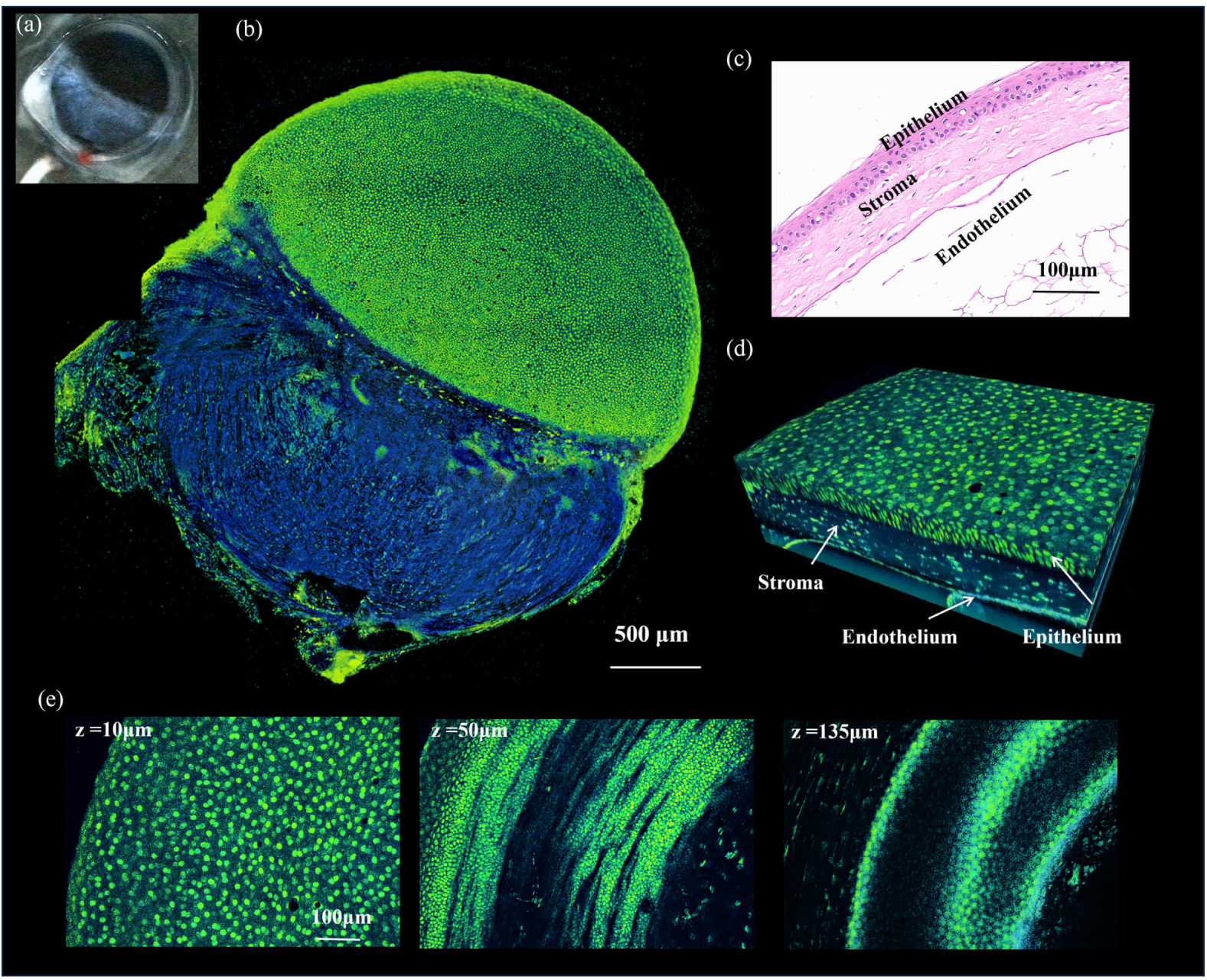
Large-area and depth-resolved D-FFOCT imaging of the fresh mouse eye. (a) Photograph of the intact fresh mouse eyeball before imaging. (b) Large-area D-FFOCT mosaic of the whole eyeball, showing the overall ocular contour and major internal structural contrast. (c) Corresponding H&E-stained histological section of the imaged corneal region, showing the epithelium, stroma, and endothelium. (d) Three-dimensional rendering of the selected corneal region, revealing its layered organization. (e) Representative en face images acquired at depths of 10, 50, and 135 µm, showing the transition from the superficial epithelial mosaic to deeper stromal structures. Scale bars: 500 µm (b); 100 µm (c–e).

The mouse eye contains distinct anatomical compartments and finely organized tissue layers, providing a useful model for evaluating large-area and depth-resolved D-FFOCT imaging. Figure 3(b) shows a large-area mosaic of the intact fresh eyeball, with an overall contour consistent with the corresponding specimen photograph (Fig. 3(a)). The image reveals the global ocular morphology and distinct contrast patterns between the anterior and posterior regions, demonstrating the ability to visualize the intact eye without fixation or tissue clearing.

To examine local microarchitecture, a corneal region was selected for histological comparison and volumetric imaging. The adjacent H&E-stained section (Fig. 3(c)) identifies the corneal epithelium, stroma, and endothelium. The three-dimensional D-FFOCT reconstruction (Fig. 3(d) and Visualization 3) reveals a corresponding layered organization, including a superficial mosaic-like texture and deeper fibrous structures. Representative en face images acquired at depths of 10, 50, and 135 µm (Fig. 3(e)) show progressive changes from a densely packed superficial pattern to elongated and curved features associated with the stromal layers. Together, these observations demonstrate that D-FFOCT can combine whole-eye morphological imaging with depth-resolved visualization of local corneal microarchitecture.

#### 2.2.3. Thymus

**Fig. 4.**
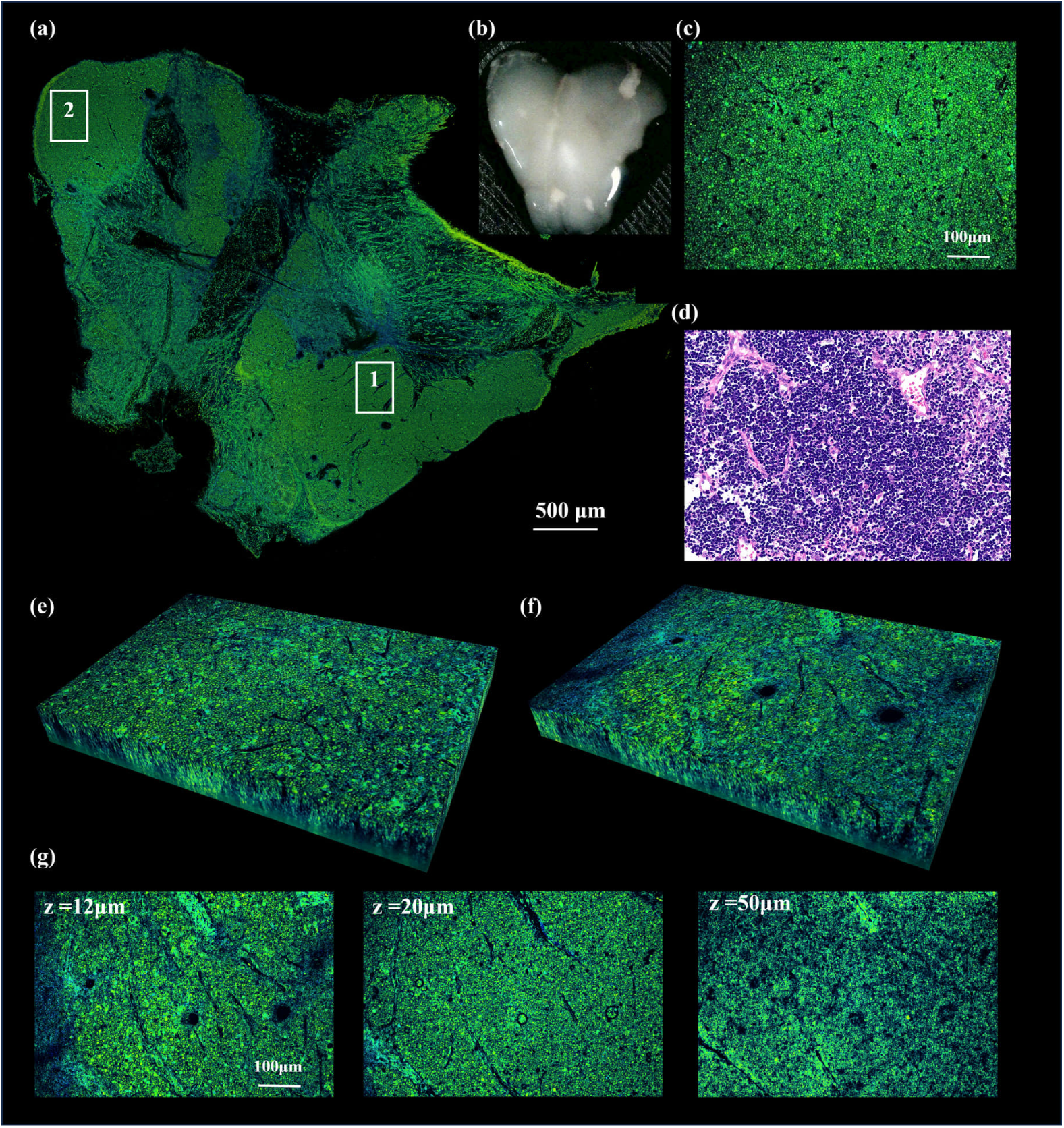
Large-area and volumetric D-FFOCT imaging of the fresh mouse thymus. (a) Large-area D-FFOCT mosaic of the freshly excised thymus, showing the overall lobulated architecture. Boxes 1 and 2 indicate representative medullary and cortical regions, respectively. (b) Photograph of the same thymus before imaging, with an overall outline consistent with that in (a). (c) Magnified D-FFOCT image of region 2, corresponding to the cortex, showing a dense and relatively uniform cellular texture. (d) Adjacent H&E-stained section of the corresponding cortical region, revealing a densely cellular lymphocyte-rich area. (e) Three-dimensional rendering of region 1, corresponding to the medulla, showing a more heterogeneous internal texture with scattered focal features. (f) Three-dimensional rendering of region 2. (g) Representative en face images of region 2 acquired at depths of 12, 20, and 50 µm, showing the persistence of the dense cortical texture and several rounded low-signal structures across depth. Scale bars: 500 µm (a); 100 µm (c–g).

The thymus is a lobulated primary lymphoid organ characterized by distinct cortical and medullary compartments. Figure 4(a) presents a large-area D-FFOCT mosaic of the freshly excised thymus, revealing a lobulated morphology consistent with the corresponding specimen photograph (Fig. 4(b)). The parenchyma exhibits regional variations in image texture, with the cortical region (region 2) appearing relatively dense and homogeneous and the medullary region (region 1) showing a more heterogeneous pattern.

The magnified D-FFOCT image of region 2 (Fig. 4(c)) reveals closely packed punctate or cell-like features. The adjacent H&E-stained section (Fig. 4(d)) confirms a densely cellular, lymphocyte-rich tissue architecture, supporting the interpretation of this region as thymic cortex. Three-dimensional imaging of region 1 (Fig. 4(e)) reveals heterogeneous medullary features, whereas region 2 (Fig. 4(f)) maintains a relatively uniform and compact texture. En face images acquired at depths of 12, 20, and 50 µm (Fig. 4(g)) further show the continuity of the cortical microarchitecture and several rounded low-signal structures morphologically consistent with small vascular profiles or tissue spaces. These results demonstrate the ability of D-FFOCT to distinguish regional structural characteristics and visualize depth-resolved microarchitecture in fresh thymic tissue.

#### 2.2.4. Inguinal lymph node

The inguinal lymph node is a secondary lymphoid organ containing distinct lymphocyte-rich compartments, medullary structures, and associated vessels. Figure 5(a) shows a large-area D-FFOCT mosaic of the freshly excised lymph node and surrounding tissue, with an overall morphology consistent with the corresponding specimen photograph (Fig. 5(b)). The parenchyma exhibits relatively dense and homogeneous textures, while localized low-signal spaces and surrounding ring-like adipose structures provide additional morphological contrast. A second compact lymphoid-appearing structure is also visible adjacent to the main specimen.

At higher magnification, region 1 (Fig. 5(c)) displays a densely packed punctate texture, consistent with the lymphocyte-rich tissue observed in the adjacent H&E-stained section. Three-dimensional reconstruction and en face images acquired at depths of 8, 20, 30, and 40 µm (Fig. 5(d)) demonstrate the continuity of the lymphoid microarchitecture. A rounded low-signal lumen-like structure remains visible across multiple depths, with localized changes in intraluminal contrast, suggesting a small vascular or sinus-like space. These results demonstrate that D-FFOCT can characterize both large-area lymphoid tissue organization and local depth-resolved microstructural features without staining or physical sectioning.

**Fig. 5.**
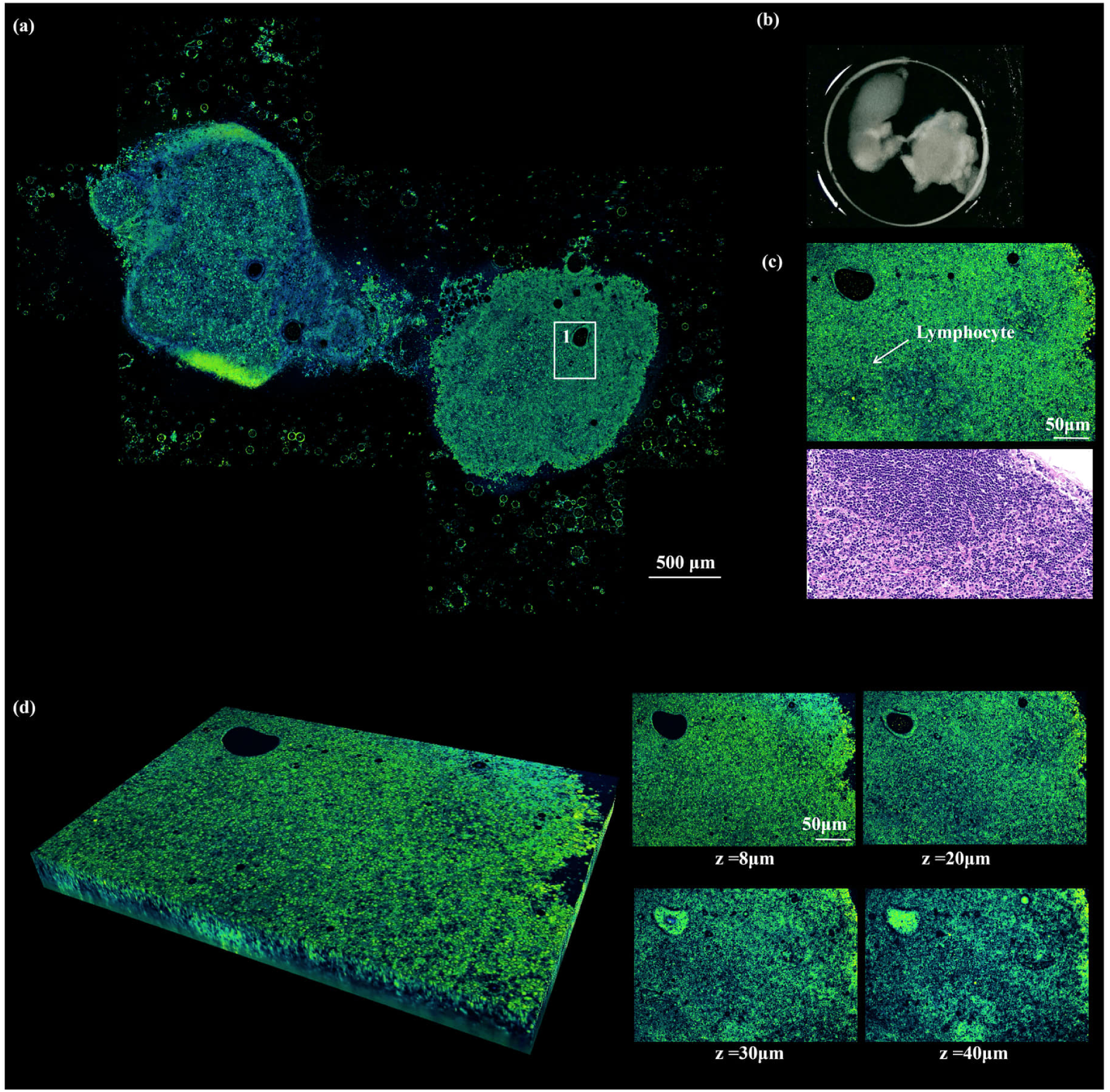
Large-area and volumetric D-FFOCT imaging of the fresh mouse inguinal lymph node. (a) Large-area D-FFOCT mosaic of the freshly excised inguinal lymph node and surrounding tissue. Box 1 indicates a representative lymphocyte-rich region selected for higher-magnification imaging. A second compact lymphoid-appearing structure is also visible within the field of view and is connected to the main specimen by a narrow tissue bridge. (b) Photograph of the same specimen before imaging, showing an overall outline consistent with that in (a). (c) Higher-magnification D-FFOCT image of region 1 and the adjacent H&E-stained section, showing a densely cellular lymphoid region. (d) Three-dimensional rendering of region 1 and representative en face images acquired at depths of 8, 20, 30, and 40 µm, showing the persistence of the dense lymphoid texture and a rounded lumen-like structure across depth. Scale bars: 500 µm (a); 50 µm (c, d).

#### 2.2.5. Liver

The liver consists of densely packed hepatocytes arranged into hepatic plates separated by sinusoidal spaces, providing characteristic cellular-scale microarchitecture. Figure 6(a) presents a large-area D-FFOCT mosaic of fresh mouse liver tissue, with the corresponding specimen photograph shown in Fig. 6(b). The image reveals a continuous parenchymal architecture containing densely distributed cellular features and fine, irregular low-signal networks, morphologically consistent with the arrangement of hepatocytes and intervening sinusoids.

The magnified image of region 1 (Fig. 6(c)) reveals polygonal hepatocyte-like profiles with relatively distinct boundaries and occasional intracellular low-signal features. The adjacent H&E-stained section shows a comparable arrangement of hepatocytes and sinusoidal spaces, supporting the morphological interpretation of the D-FFOCT image. Three-dimensional reconstruction (Fig. 6(d) and Visualization 4) and representative en face images at depths of 5, 20, 80, and 120 µm further illustrate the depth-dependent appearance of hepatic microarchitecture. Individual cellular profiles are most clearly delineated at shallow depths, while deeper images exhibit reduced cellular contrast but retain larger-scale parenchymal features. These findings demonstrate the ability of D-FFOCT to combine large-area hepatic tissue visualization with cellular-scale morphological characterization and local volumetric imaging.

**Fig. 6.**
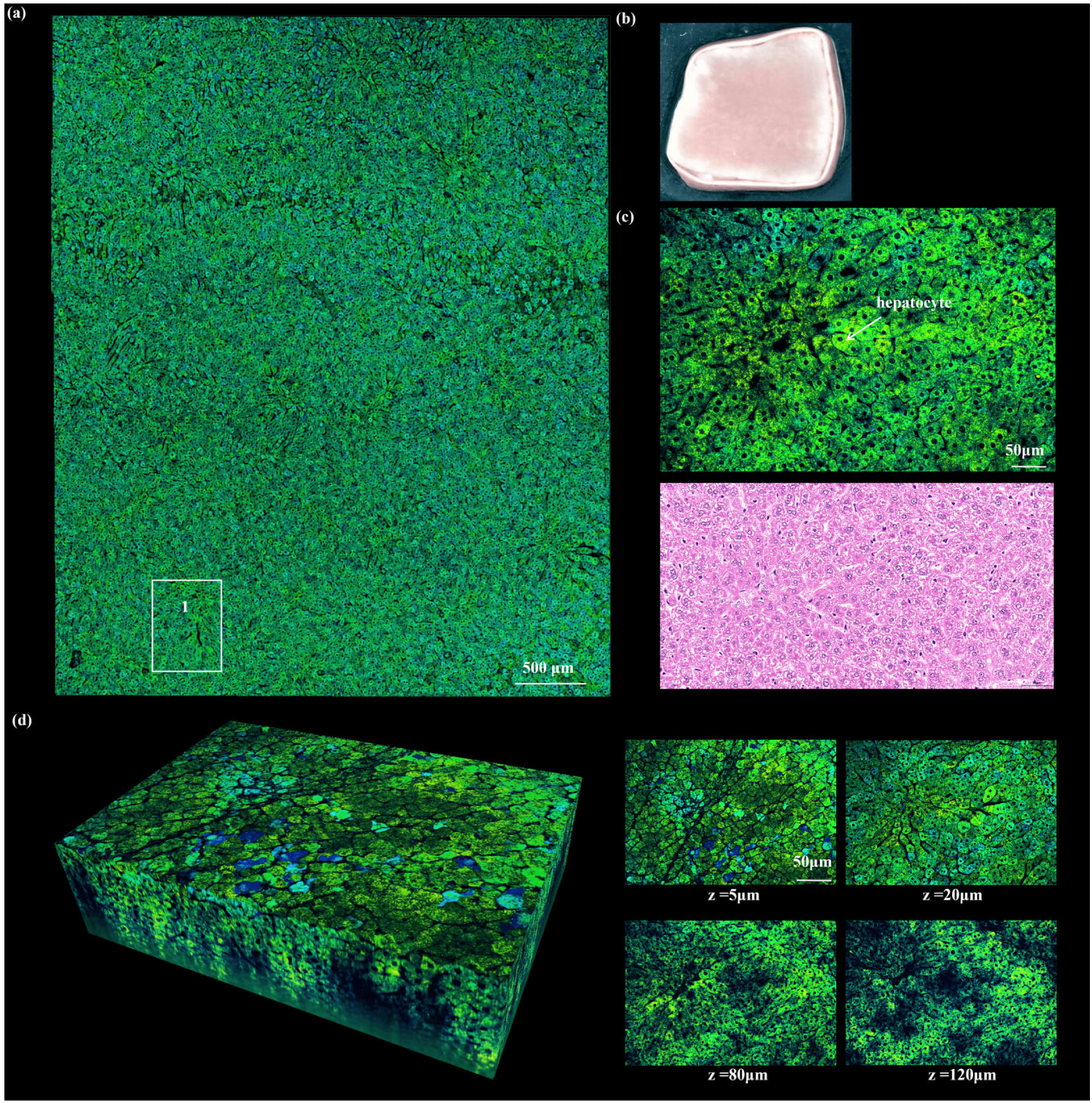
Large-area and volumetric D-FFOCT imaging of fresh mouse liver tissue. (a) Large-area D-FFOCT mosaic of a freshly excised mouse liver section, showing the densely organized hepatic parenchyma and spatially distributed cellular features. Box 1 indicates a representative parenchymal region selected for higher-magnification and volumetric imaging. (b) Photograph of the corresponding fresh liver specimen before imaging. (c) Magnified D-FFOCT image of region 1 (top) and an adjacent H&E-stained histological section (bottom), showing hepatocyte-like cellular profiles and the surrounding sinusoid-like spaces. (d) Three-dimensional reconstruction of region 1 (left) and representative en face images acquired at depths of 5, 20, 80, and 120 µm (right), illustrating the depth-dependent appearance of hepatic cellular structures and changes in image contrast. Scale bars: 500 µm (a); 50 µm (c, d).

#### 2.2.6. Pancreas

The pancreas is a mixed exocrine–endocrine gland composed predominantly of densely packed serous acini, with scattered endocrine islets and connective tissue septa. Figure 7(a) presents a large-area D-FFOCT mosaic of freshly excised mouse pancreatic tissue, showing a lobulated architecture consistent with the corresponding specimen photograph (Fig. 7(b)). The exocrine parenchyma exhibits a dense, finely granular texture, while localized linear and band-like structures delineate different tissue regions, morphologically consistent with interlobular connective tissue.

**Fig. 7.**
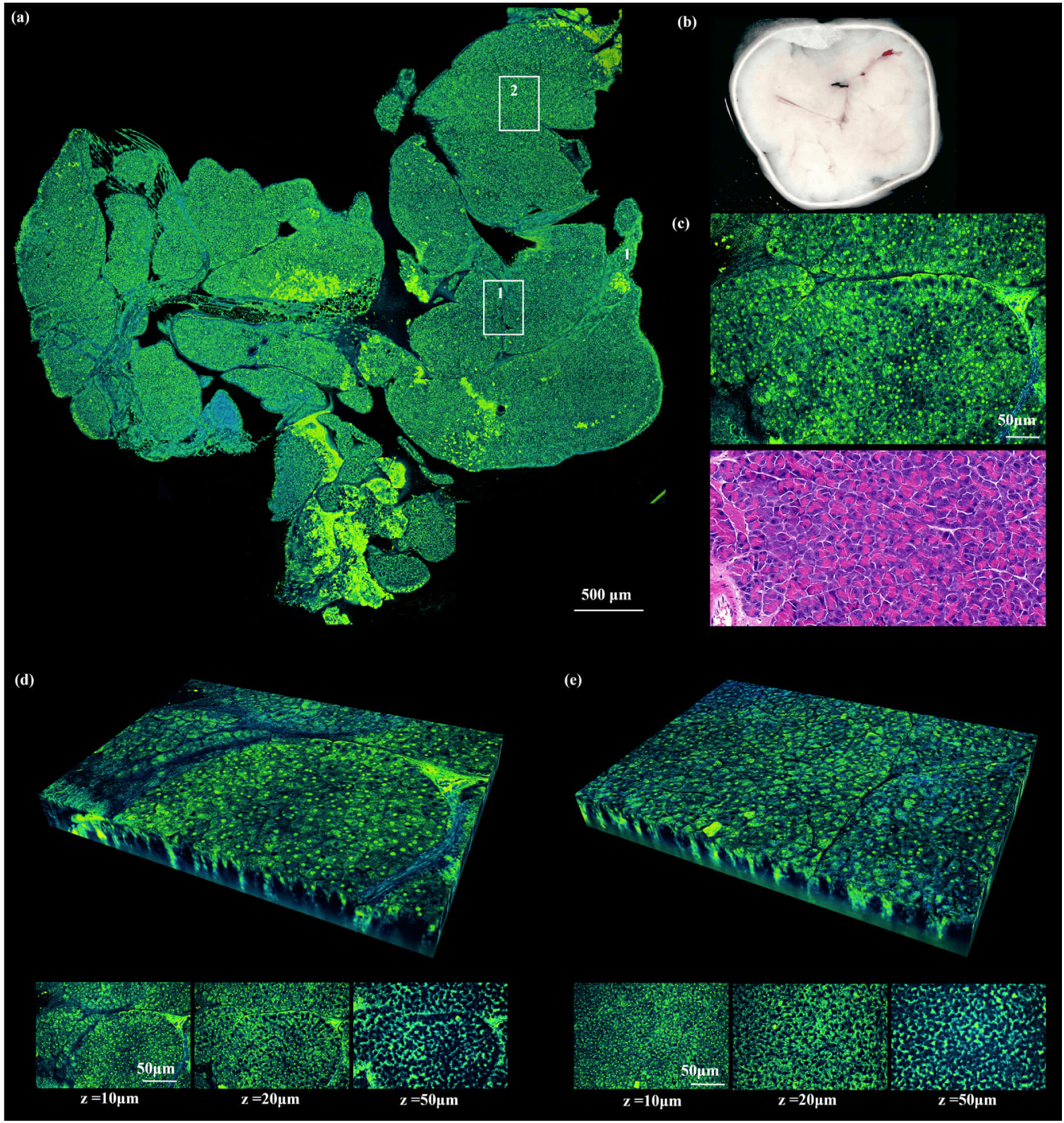
Large-area and volumetric D-FFOCT imaging of fresh mouse pancreatic tissue. (a) Large-area D-FFOCT mosaic of the freshly excised pancreas. Boxes 1 and 2 indicate representative regions selected for detailed imaging. (b) Photograph of the fresh pancreatic specimen before imaging. (c) Magnified D-FFOCT image of region 1 (top) and an adjacent H&E-stained histological section (bottom), showing a septum-like structure surrounded by densely packed acinar tissue. (d) Three-dimensional reconstruction of region 1 and representative en face images acquired at depths of 10, 20, and 50 µm, revealing the septum-associated structure and adjacent granular parenchyma. (e) Three-dimensional reconstruction of region 2 and corresponding en face images at depths of 10, 20, and 50 µm, showing relatively homogeneous granular tissue with scattered low-signal spaces. Scale bars: 500 µm (a); 50 µm (c–e).

At higher magnification, region 1 (Fig. 7(c)) reveals a continuous septum-like structure surrounded by densely packed granular tissue. The adjacent H&E-stained section shows a comparable arrangement of connective tissue and serous acini, supporting the morphological interpretation of the D-FFOCT image. Three-dimensional imaging of region 1 (Fig. 7(d)) further demonstrates the continuity of the septum-associated structure across depths of 10, 20, and 50 µm. In contrast, region 2 (Fig. 7(e)) exhibits a relatively homogeneous granular texture throughout the imaged volume, consistent with acinar-rich exocrine parenchyma, with scattered fine low-signal spaces. Together, these results demonstrate the ability of D-FFOCT to resolve both large-area pancreatic organization and local depth-dependent microarchitecture in fresh, unstained tissue.

#### 2.2.7. Duodenum

**Fig. 8.**
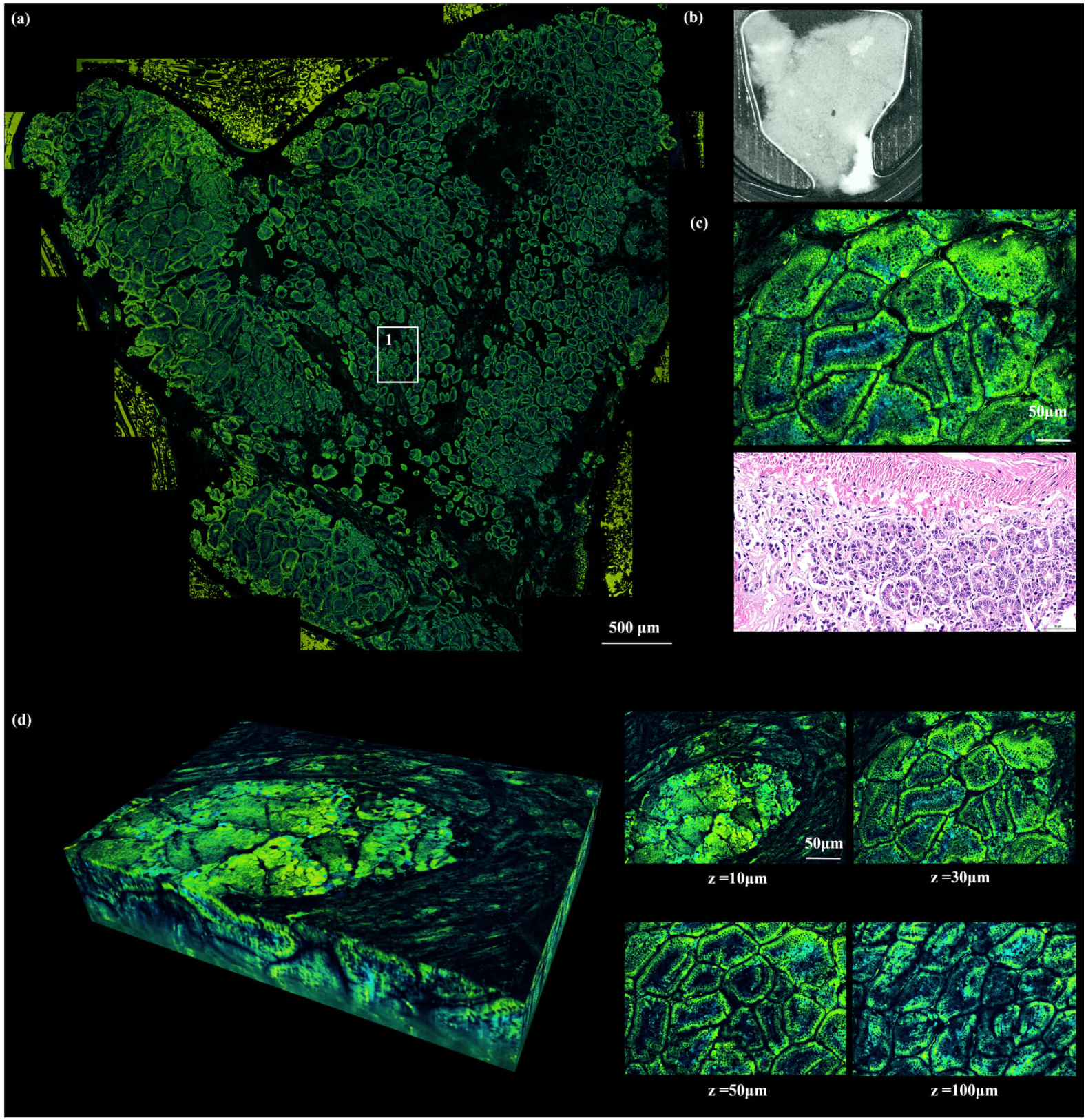
Large-area and volumetric D-FFOCT imaging of fresh mouse duodenal tissue. (a) Large-area D-FFOCT mosaic of the freshly opened duodenal mucosal surface. Box 1 indicates the representative region selected for detailed imaging. (b) Photograph of the corresponding fresh duodenal specimen before imaging. (c) Magnified D-FFOCT image of region 1 (top) and an adjacent H&E-stained histological section (bottom), showing villus profiles with bright peripheral contours and relatively lower-signal central regions. (d) Three-dimensional reconstruction of region 1 and representative en face images acquired at depths of 10, 30, 50, and 100 µm, showing the persistence of villus-associated structures and their depth-dependent transition toward deeper mucosal architecture. Scale bars: 500 µm (a); 50 µm (c, d).

The duodenum contains characteristic mucosal villi and crypt structures that provide distinctive surface and depth-resolved microarchitectural features for label-free imaging. Figure 8(a) shows a large-area D-FFOCT mosaic of the freshly opened duodenal mucosal surface, with an overall morphology consistent with the corresponding specimen photograph (Fig. 8(b)). The image reveals densely distributed villus-like structures with clear luminal contours, allowing the mucosal architecture to be visualized over a relatively large field of view.

At higher magnification, region 1 (Fig. 8(c)) shows multiple villus profiles outlined by bright ring-like or band-like contours surrounding relatively lower-signal central regions. Low-signal spaces between adjacent villi are also visible. The adjacent H&E-stained section shows comparable villus morphology and glandular organization, supporting the interpretation that the D-FFOCT image captures the villus epithelium and core mucosal architecture.

Three-dimensional imaging of region 1 (Fig. 8(d)) further demonstrates the depth-dependent organization of the duodenal mucosa. Representative en face images acquired at depths of 10, 30, 50, and 100 µm show that the villus-associated ring-like structures remain recognizable across depth, while the spaces between villi progressively narrow and transition toward deeper mucosal structures. Together, these results demonstrate that D-FFOCT can resolve both large-area mucosal organization and local depth-resolved villus microarchitecture in fresh, unstained duodenal tissue.

#### 2.2.8. Kidney

The kidney contains distinct cortical and medullary compartments, with characteristic tubular and glomerular microarchitecture that provides clear structural contrast for label-free imaging. Figure 9(a) shows a large-area D-FFOCT mosaic of a freshly excised mouse kidney section, with an overall morphology consistent with the corresponding specimen photograph (Fig. 9(b)). The image reveals a clear corticomedullary organization. Region 1 exhibits a predominantly parallel, stripe-like pattern, consistent with aligned tubular structures, whereas region 2 shows a more heterogeneous arrangement of rounded and irregular tubular profiles, consistent with cortical microarchitecture. Region 3 was selected for higher-magnification imaging and comparison with the corresponding histological section.

At higher magnification, region 3 (Fig. 9(c)) shows multiple tubular profiles with relatively bright walls and lower-signal luminal spaces, together with scattered rounded structures morphologically consistent with glomeruli. The adjacent H&E-stained section shows a comparable distribution of renal tubules and renal corpuscle-like structures, supporting the morphological interpretation of the D-FFOCT image.

Three-dimensional imaging further highlights the depth-resolved organization of these renal structures. Region 1 (Fig. 9(d)) preserves a predominantly parallel, stripe-like pattern across depths of 10, 30, and 50 µm, consistent with the ordered arrangement of tubular structures. In contrast, region 2 (Fig. 9(e)) shows a more heterogeneous pattern of rounded and irregular tubular profiles across the same depth range, consistent with cortical microarchitecture. Together, these results demonstrate that D-FFOCT can visualize distinct renal microarchitectural patterns and their spatial organization across imaging depths in fresh, unstained kidney tissue.

**Fig. 9.**
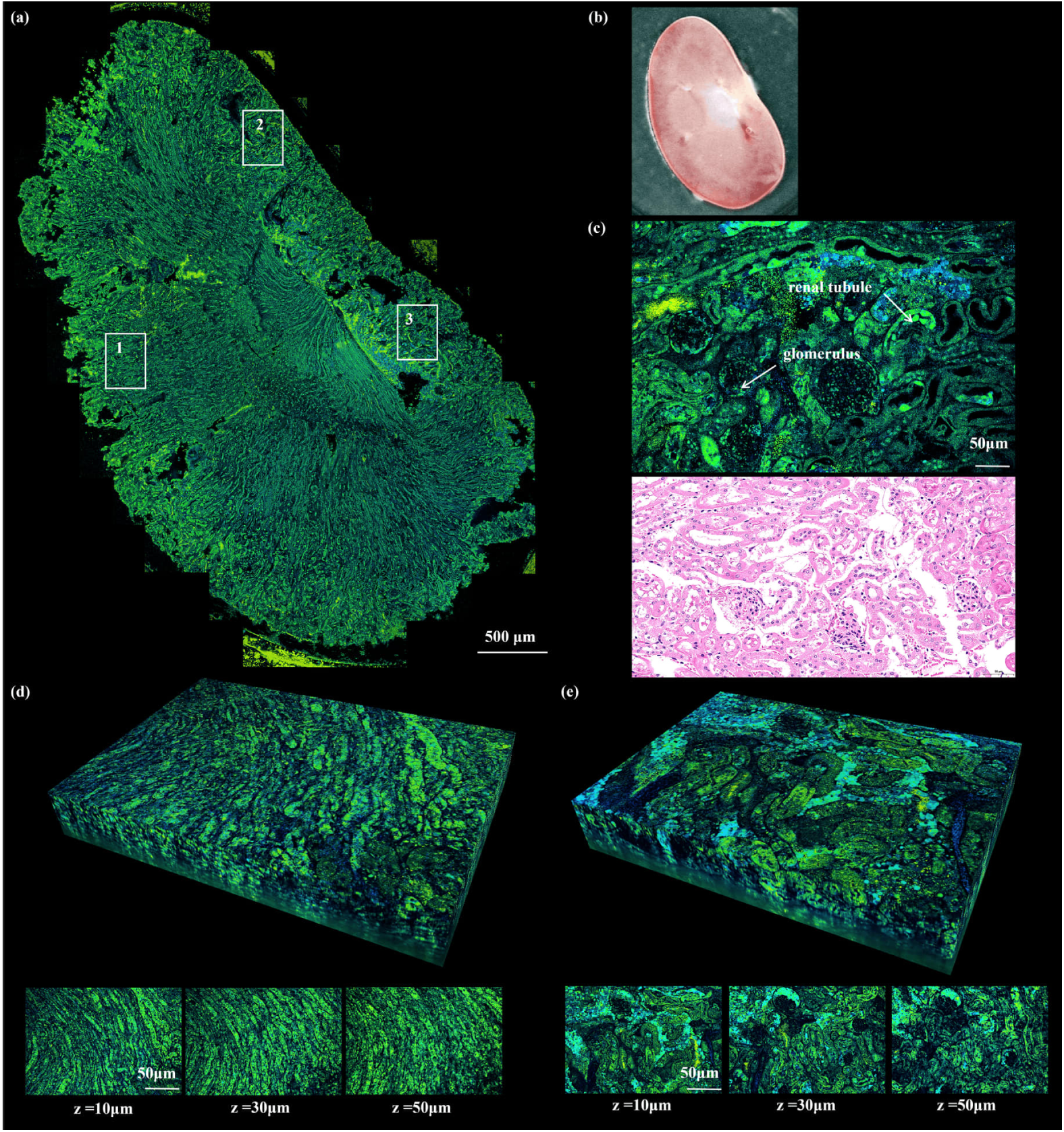
Large-area and volumetric D-FFOCT imaging of fresh mouse kidney tissue. (a) Large-area D-FFOCT mosaic of a freshly excised mouse kidney section. Boxes 1–3 indicate representative regions selected for detailed imaging. (b) Photograph of the corresponding fresh kidney specimen before imaging. (c) Magnified D-FFOCT image of region 3 (top) and an adjacent H&E-stained histological section (bottom), showing tubular structures with relatively bright walls and lower-signal lumina together with scattered rounded glomerulus-like structures. (d) Three-dimensional reconstruction of region 1 and representative en face images acquired at depths of 10, 30, and 50 µm, showing predominantly parallel, stripe-like tubular structures. (e) Three-dimensional reconstruction of region 2 and corresponding en face images at depths of 10, 30, and 50 µm, showing a heterogeneous arrangement of rounded and irregular tubular profiles consistent with cortical microarchitecture. Scale bars: 500 µm (a); 50 µm (c–e).

#### 2.2.9. Ovary

The ovary contains heterogeneous structural domains, including the outer cortex with follicles at different developmental stages and central stromal or medullary regions, making it well suited for label-free structural imaging. Figure 10(a) shows a large-area D-FFOCT mosaic of a freshly excised mouse ovary, with an overall morphology consistent with the corresponding specimen photograph (Fig. 10(b)). The ovarian surface is outlined by a relatively bright peripheral rim, while the parenchyma contains both compact solid regions and multiple rounded ring-like structures. Region 1 appears as a dense, high-signal solid structure with a central lower-signal area, morphologically consistent with a corpus luteum-like region, whereas region 2 contains multiple rounded follicle-like structures distributed within the cortex.

**Fig. 10.**
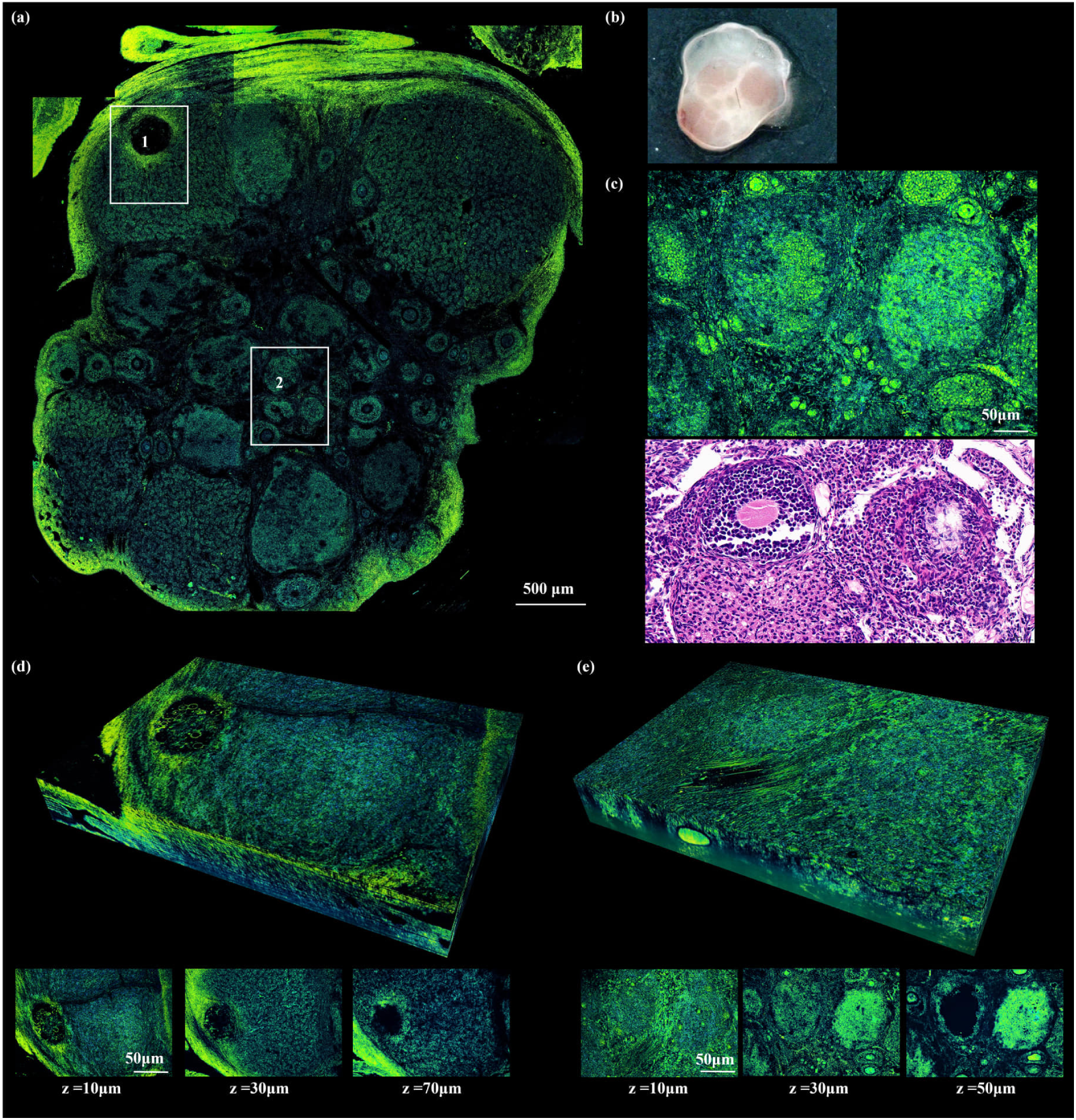
Large-area and volumetric D-FFOCT imaging of fresh mouse ovarian tissue. (a) Large-area D-FFOCT mosaic of a freshly excised mouse ovary. Boxes 1 and 2 indicate representative regions selected for detailed imaging. (b) Photograph of the corresponding fresh ovarian specimen before imaging. (c) Magnified D-FFOCT image of region 2 (top) and an adjacent H&E-stained histological section (bottom), showing multiple follicle-like structures with moderately to strongly scattering outer rims and relatively lower-signal central regions. (d) Three-dimensional reconstruction of region 1 and representative en face images acquired at depths of 10, 30, and 70 µm, showing a compact high-signal corpus luteum-like structure with a persistent central lower-signal region. (e) Three-dimensional reconstruction of region 2 and corresponding en face images acquired at depths of 10, 30, and 50 µm, showing multiple rounded follicle-like profiles across depth. Scale bars: 500 µm (a); 50 µm (c–e).

At higher magnification, region 2 (Fig. 10(c)) reveals several rounded structures composed of moderately to strongly scattering outer rims surrounding relatively lower-signal central regions. The adjacent H&E-stained section shows a comparable distribution of developing follicles with layered follicular walls and central follicular spaces, supporting the interpretation of these structures as follicles at different developmental stages.

Three-dimensional imaging further highlights the depth-resolved organization of ovarian structures. Region 1 (Fig. 10(d)) remains as a compact high-signal mass across depths of 10, 30, and 70 µm, with a persistent central lower-signal region. In contrast, region 2 (Fig. 10(e)) preserves multiple rounded ring-like follicular profiles across depths of 10, 30, and 50 µm, with outer cellular rims and lower-signal centers remaining recognizable throughout the imaged volume. Together, these results demonstrate that D-FFOCT can resolve both large-area ovarian organization and local depth-resolved follicular and luteal microarchitecture in fresh, unstained tissue.

#### 2.2.10. Testis

**Fig. 11.**
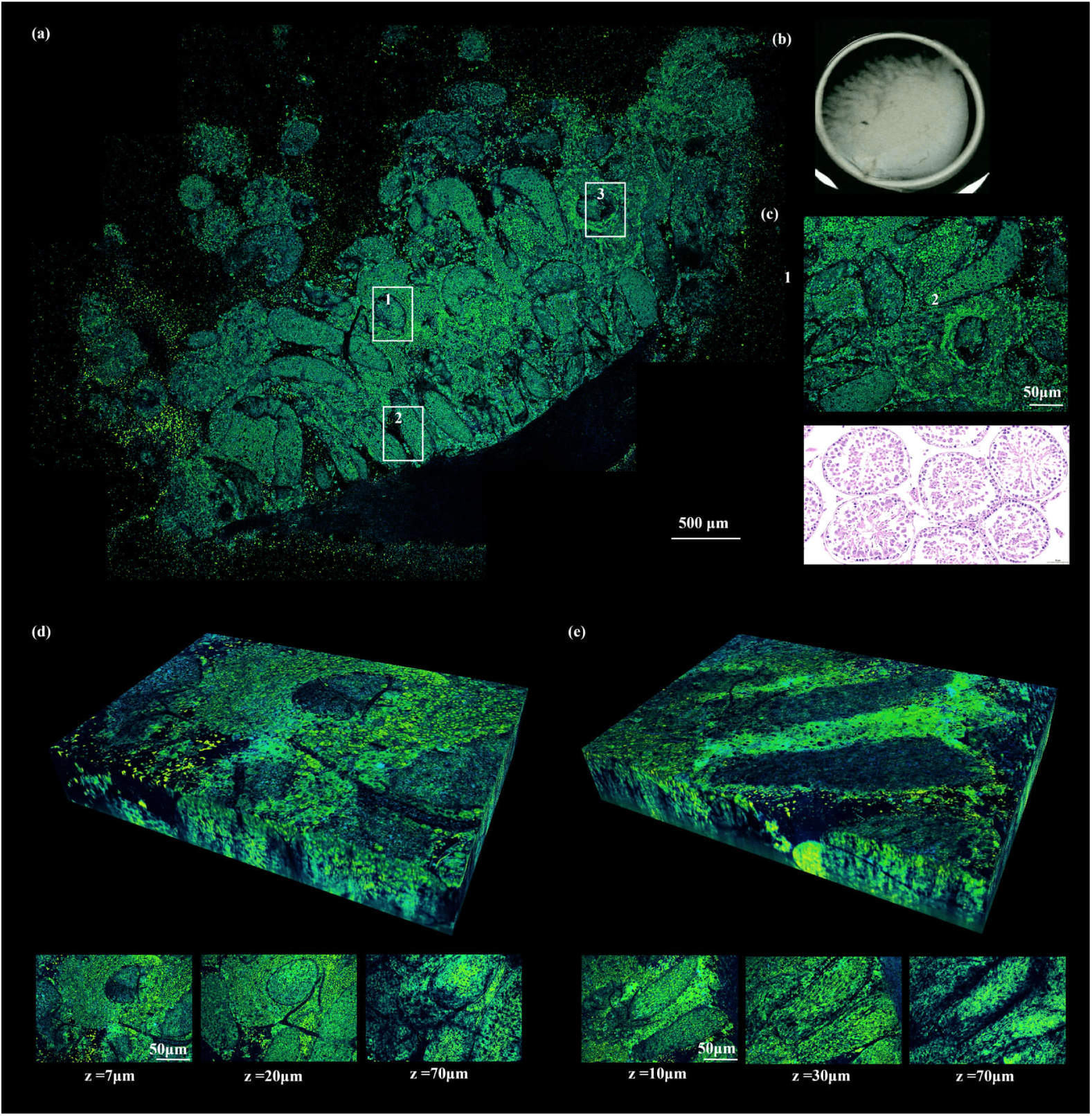
Large-area and volumetric D-FFOCT imaging of fresh mouse testicular tissue. (a) Large-area D-FFOCT mosaic of a freshly excised mouse testis section, showing densely distributed seminiferous tubules with rounded and elongated profiles. Boxes 1–3 indicate representative regions within the tissue. (b) Photograph of the corresponding fresh testis specimen before imaging. (c) Magnified D-FFOCT image of region 1 (top) and an adjacent H&E-stained histological section (bottom), showing seminiferous tubule profiles with distinct peripheral epithelial layers and central luminal regions. (d) Three-dimensional reconstruction of region 1 and representative en face images acquired at depths of 7, 20, and 70 µm, showing rounded seminiferous tubule profiles and surrounding interstitial tissue. (e) Three-dimensional reconstruction of region 2 and corresponding en face images acquired at depths of 10, 30, and 70 µm, showing elongated seminiferous tubule structures and their depth-dependent organization. Scale bars: 500 µm (a); 50 µm (c–e).

The testis consists of densely packed seminiferous tubules surrounded by interstitial tissue, providing a characteristic tubular architecture for label-free imaging. Figure 11(a) presents a large-area D-FFOCT mosaic of a freshly excised mouse testis section, showing numerous rounded and elongated seminiferous tubule profiles consistent with the corresponding specimen photograph (Fig. 11(b)). The tubules exhibit relatively well-defined peripheral boundaries surrounding lower-signal central regions, while the intervening interstitial tissue forms irregular branching patterns.

At higher magnification, region 1 (Fig. 11(c)) reveals seminiferous tubule profiles with distinct peripheral cellular layers and central luminal regions. The adjacent H&E-stained section shows comparable seminiferous epithelial organization and intertubular spaces, supporting the morphological interpretation of the D-FFOCT image.

Three-dimensional imaging further illustrates the depth-resolved arrangement of these tubular structures. Region 1 (Fig. 11(d)) shows rounded tubule profiles and surrounding interstitial regions across depths of 7, 20, and 70 µm. Region 2 (Fig. 11(e)) contains more elongated and closely arranged tubule profiles, with their characteristic boundaries remaining recognizable at depths of 10, 30, and 70 µm. Changes in tubule shape across depth reflect their curved spatial trajectories and different sectioning orientations. Together, these results demonstrate that D-FFOCT can visualize both the large-area organization and local three-dimensional microarchitecture of seminiferous tubules in fresh, unstained testicular tissue.

Collectively, these results demonstrate the broad applicability of D-FFOCT for label-free imaging of diverse fresh mouse organs. Across ten organs, the platform reveals distinct tissue-specific microarchitectures, ranging from the layered organization of the brain and cornea to the cellular and glandular structures of the liver, pancreas, kidney, and reproductive organs. Large-area mosaics provide spatial context for identifying regional tissue organization, while local histological comparisons and three-dimensional reconstructions further support the interpretation of microscopic structures and their depth-dependent arrangement. Importantly, these observations demonstrate that D-FFOCT can capture complementary morphological and endogenous dynamic contrast across tissues with substantially different cellular compositions and structural organizations. Together, these findings establish a systematic volumetric reference of normal mouse organs and provide a foundation for subsequent cross-organ quantitative analysis of tissue dynamics.

### 2.3 Quantitative comparison of dynamic characteristics across mouse organs

Beyond morphological characterization, D-FFOCT provides quantitative information on endogenous tissue dynamics through temporal fluctuations in interferometric signals. To investigate organ-dependent differences, we analyzed the mean dynamic frequency (Hmean) and mean dynamic intensity (Vmean) across 120 imaging locations from nine mouse organs, including the brain, eyeball, thymus, inguinal lymph node, liver, pancreas, duodenum, kidney, and testis (9–17 locations per organ). Hmean characterizes the temporal frequency distribution of signal fluctuations, whereas Vmean describes their magnitude, providing complementary measures of tissue dynamics.

As shown in Fig. 12(a), Hmean exhibited pronounced differences among organs (one-way ANOVA, p < 0.0001). The pancreas showed the highest mean frequency (77.49 ± 5.78 Hz), followed by the kidney (75.49 ± 6.06 Hz) and brain (74.25 ± 6.18 Hz), whereas the liver exhibited the lowest value (52.02 ± 3.70 Hz). The remaining organs displayed intermediate values, with the eyeball and inguinal lymph node generally showing higher frequencies than the thymus, testis, and duodenum. These differences indicate that the temporal characteristics of endogenous interferometric fluctuations vary substantially across tissue types.

In contrast, Vmean exhibited a markedly different organ-dependent distribution (Fig. 12(b)). The liver showed the highest dynamic intensity (45.08 ± 8.08 a.u.), followed by the testis (33.14 ± 2.95 a.u.), whereas the pancreas exhibited the lowest value (15.91 ± 1.29 a.u.). Other organs displayed intermediate dynamic intensities, generally ranging from approximately 21 to 28 a.u. Notably, the liver and pancreas occupied opposite extremes in the two measurements: the liver exhibited relatively low-frequency but high-intensity fluctuations, while the pancreas showed high-frequency but low-intensity fluctuations. This contrasting behavior suggests that the characteristic frequency and magnitude of interferometric fluctuations capture different aspects of endogenous tissue dynamics, which may reflect differences in cellular organization and intracellular motion.

Together, these results reveal distinct organ-dependent dynamic signatures and demonstrate the value of combining frequency- and intensity-based measurements for quantitative tissue characterization. Although the underlying biological mechanisms require further investigation, the observed differences provide a comparative reference for understanding endogenous dynamic contrast across diverse fresh mouse organs.

**Fig. 12.**
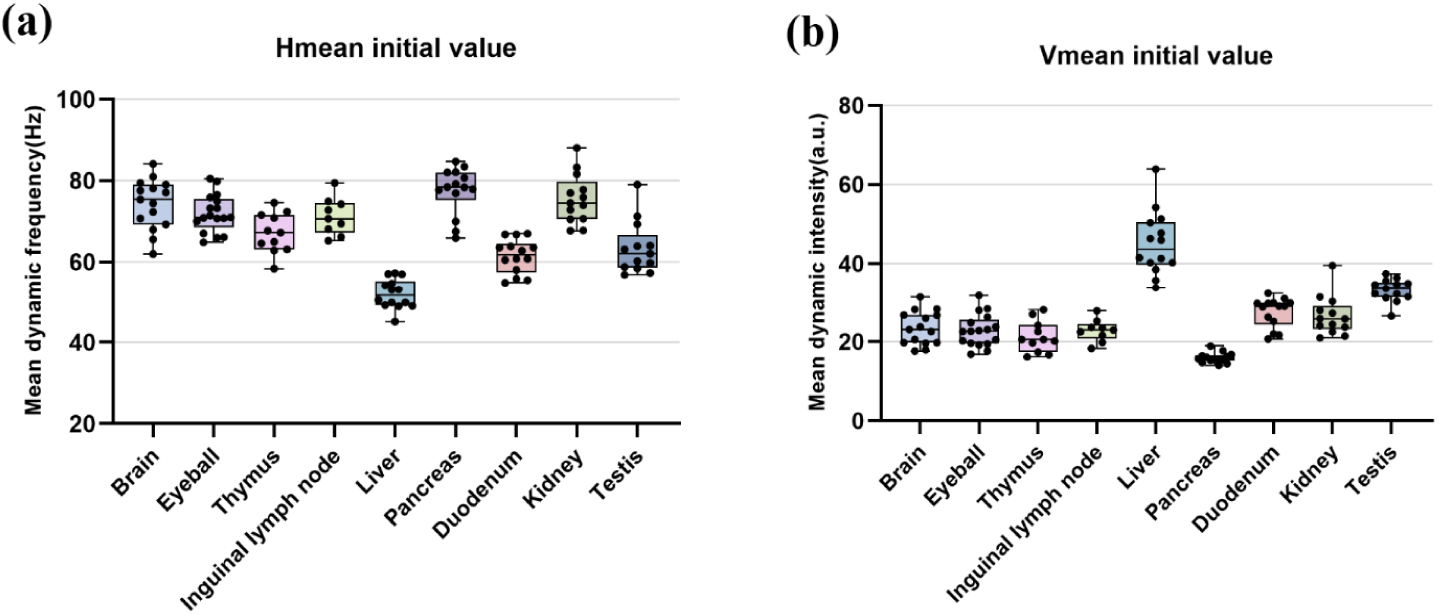
Quantitative comparison of D-FFOCT dynamic characteristics across nine fresh mouse organs. (a) Box plots showing the mean dynamic frequency (Hmean, Hz) measured in the brain, eyeball, thymus, inguinal lymph node, liver, pancreas, duodenum, kidney, and testis. (b) Corresponding distributions of mean dynamic intensity (Vmean, a.u.) across the same organs. Each black dot represents one imaging location (120 locations in total, with 9–17 locations per organ). Boxes indicate the interquartile range and median, and whiskers represent the range of non-outlier values.

### 2.4. Longitudinal evolution of endogenous dynamics in freshly excised tissues

Beyond cross-organ comparisons, we investigated the temporal evolution of endogenous dynamic contrast in freshly excised mouse brain, liver, and duodenal tissues using longitudinal D-FFOCT imaging (Figs. 13–15). The observation periods were 30, 20, and 15 h, respectively, with repeated measurements at selected time points. Between acquisitions, the LED illumination was automatically switched off to minimize unnecessary light exposure. At each time point, we acquired 1,024 temporal frames and extracted the mean dynamic frequency (Hmean) and mean dynamic intensity (Vmean) from 20 tracked cellular or local tissue regions per organ. These measurements enabled the characterization of organ-dependent changes in interferometric fluctuations alongside their corresponding morphological features.

#### 2.4.1 Brain: dentate gyrus

Figures 13(a)–13(f) show the longitudinal evolution of D-FFOCT dynamic contrast in freshly excised mouse dentate gyrus tissue over 30 h (Visualization 5). At baseline, the densely packed cellular region exhibits pronounced intracellular dynamic contrast, with distinguishable cell-like profiles and heterogeneous internal textures. Notably, substantial changes become apparent during the early observation period: the intracellular dynamic signals weaken rapidly, and many cellular regions become less distinguishable from their surroundings, leaving predominantly residual cellular outlines and the broader layered architecture. During subsequent hours, the overall tissue organization remains recognizable, whereas the intracellular textures become increasingly homogeneous. By 30 h, the fine cellular dynamic contrast is substantially diminished, although portions of the original structural framework remain visible.

**Fig. 13.**
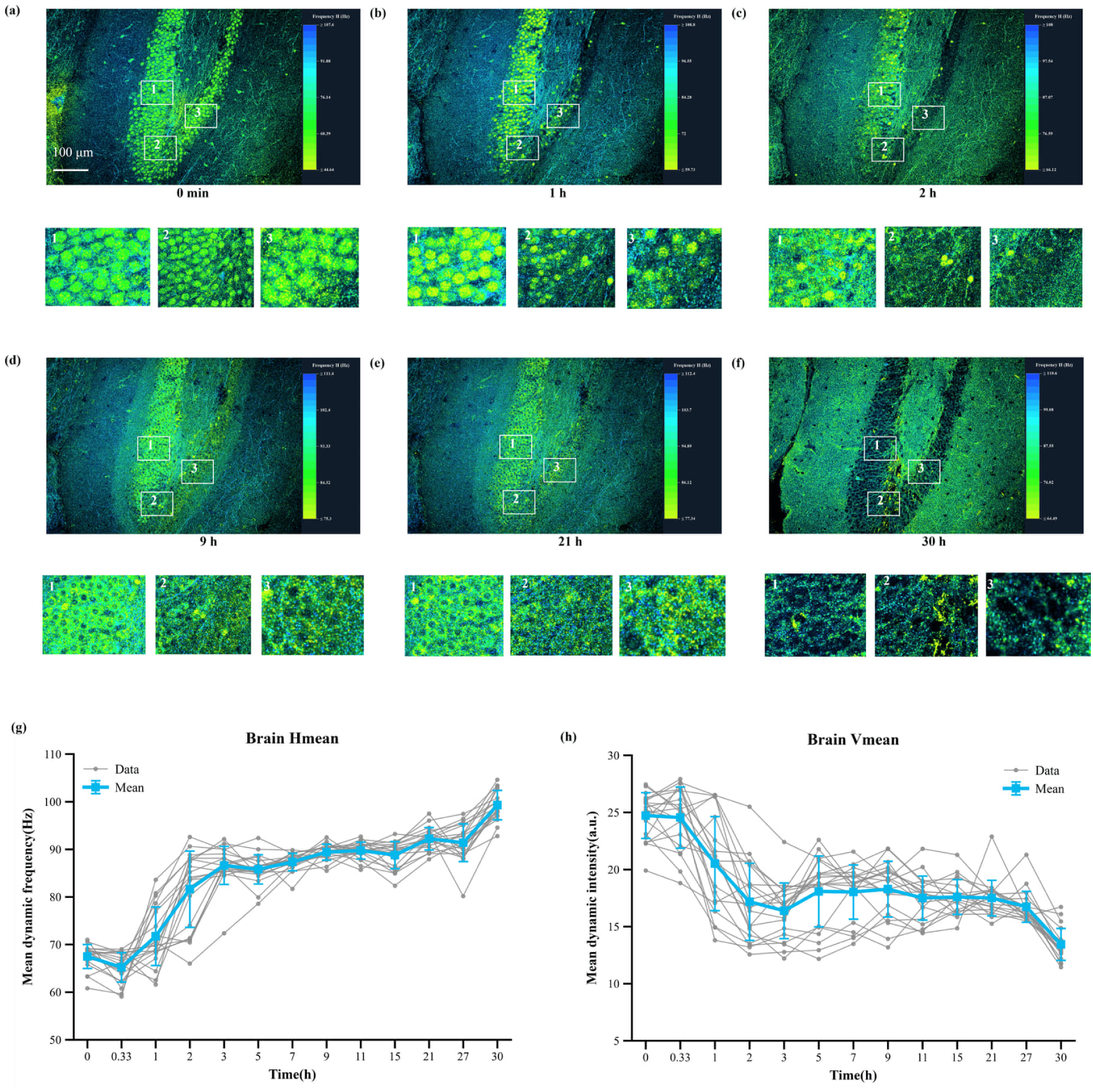
Longitudinal changes in endogenous D-FFOCT dynamics of freshly excised mouse dentate gyrus tissue. (a)–(f) Representative dynamic frequency maps of the same tissue region acquired at 0, 1, 2, 9, 21, and 30 h after excision, respectively. Three numbered regions of interest are indicated in each image, with corresponding magnified views shown below to illustrate local morphological and dynamic contrast changes. (g) Temporal evolution of mean dynamic frequency (Hmean, Hz) measured at 13 time points over 30 h. (h) Corresponding changes in mean dynamic intensity (Vmean, a.u.). Gray lines represent individual tracked cellular or local regions (n = 20), and blue markers and error bars indicate the mean ± standard deviation. Time points are displayed at equally spaced positions along the horizontal axis. Scale bar: 100 µm.

Quantitative measurements are consistent with this rapid early change. Vmean decreases from 24.73 ± 2.00 a.u. at baseline to approximately 16.38 a.u. within the first 3 h, corresponding to a reduction of approximately 34%, followed by a prolonged period of relatively stable low values and a further decline to 13.44 ± 1.40 a.u. at 30 h (Fig. 13(h)). Meanwhile, Hmean increases from 67.49 ± 2.52 Hz to approximately 86.62 Hz during the first 3 h, remains relatively stable over the intermediate period, and reaches 99.29 ± 3.09 Hz at 30 h (Fig. 13(g)). Together, these results indicate that the loss of intracellular dynamic contrast occurs rapidly after excision, while cellular outlines and the larger-scale tissue architecture remain partially preserved, revealing a temporal distinction between endogenous dynamic contrast and recognizable tissue morphology.

#### 2.4.2. Liver

**Fig. 14.**
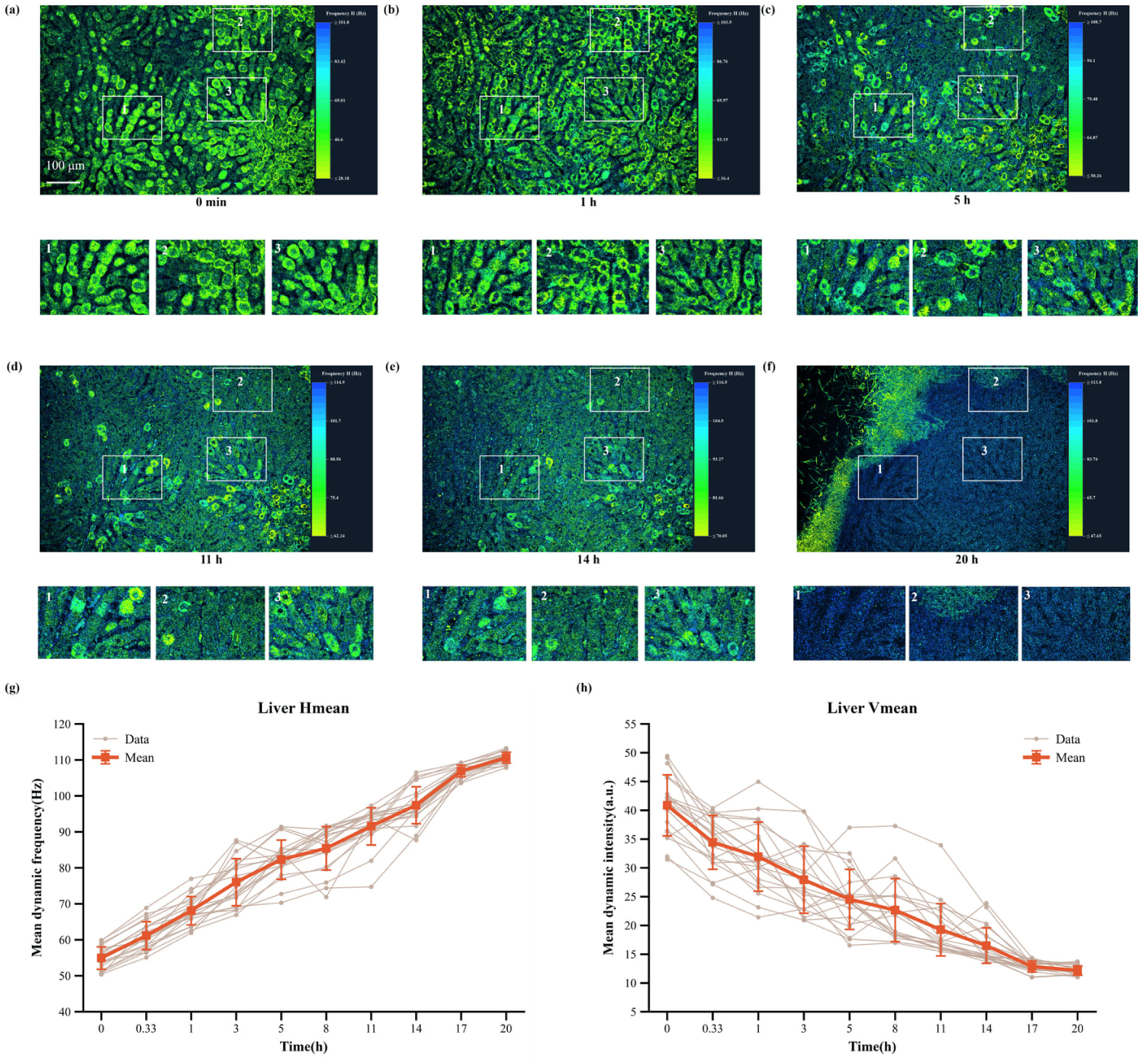
Longitudinal changes in endogenous D-FFOCT dynamics of freshly excised mouse liver tissue. (a)–(f) Representative dynamic frequency maps of the same liver region acquired at 0, 1, 5, 11, 14, and 20 h after excision, respectively. Magnified views of three numbered regions of interest are shown below each image, illustrating progressive changes in hepatocyte-scale morphology and endogenous dynamic contrast. (g) Temporal evolution of mean dynamic frequency (Hmean, Hz) over 20 h. (h) Corresponding changes in mean dynamic intensity (Vmean, a.u.). Gray lines represent individual tracked cellular or local regions (n = 20), and orange-red markers and error bars indicate the mean ± standard deviation. Measurements were obtained at 10 time points, displayed at equally spaced positions along the horizontal axis. Scale bar: 100 µm.

Figures 14(a)–14(f) illustrate the longitudinal evolution of D-FFOCT dynamic contrast in freshly excised mouse liver tissue over 20 h (Visualization 6). At baseline, the hepatic parenchyma exhibits pronounced intracellular dynamic contrast, with numerous hepatocyte-like profiles and heterogeneous granular textures clearly distinguishable. Unlike the rapid early loss of intracellular contrast observed in the brain, liver tissue retains recognizable cellular profiles and substantial local dynamic contrast during the initial observation period. As time progresses, the intracellular signal becomes increasingly heterogeneous and gradually weakens across the tissue. By 11–14 h, many cellular regions exhibit reduced dynamic contrast, although portions of the original cellular organization remain identifiable. At 20 h, most of the imaged parenchyma displays markedly diminished intracellular contrast, with only scattered structural features and localized residual signals remaining visible. This progressive reduction in dynamic contrast occurs without an equally abrupt disappearance of the larger-scale tissue organization.

Quantitative analysis further reveals a sustained evolution of hepatic dynamic characteristics. Hmean increases from 54.93 ± 3.13 Hz at baseline to approximately 110.60 Hz at 20 h (Fig. 14(g)), representing an overall increase of approximately 101%. In contrast, Vmean decreases progressively from 40.86 to 12.17 a.u. (Fig. 14(h)), corresponding to a reduction of approximately 70%. Unlike the brain, where dynamic intensity decreases rapidly during the first few hours and subsequently enters a prolonged relatively stable phase, liver Vmean continues to decline throughout the observation period, with a more pronounced reduction during the initial hours followed by a gradual decrease. Notably, the progressive loss of intracellular dynamic contrast observed in the images is consistent with the sustained reduction in Vmean, while the increasing Hmean indicates a concurrent shift in the temporal frequency distribution of the remaining fluctuations. Together, these findings suggest that freshly excised liver tissue undergoes a gradual and sustained alteration in endogenous dynamic contrast, distinct from the rapid early transition and prolonged stabilization observed in brain tissue.

Unlike the brain, liver Hmean increased continuously throughout the observation period, rising from 54.93 ± 3.13 Hz at baseline to approximately 110.60 Hz at 20 h (Fig. 14(g)). Over the same interval, Vmean decreased progressively from 40.86 to 12.17 a.u., representing a reduction of approximately 70% (Fig. 14(h)). The strongest changes in dynamic intensity occurred during the early observation period, followed by a more gradual decline. Notably, the substantial deterioration of fine morphological contrast at later time points coincided with elevated Hmean and markedly reduced Vmean. These results reveal a sustained temporal shift in hepatic dynamic characteristics following excision, accompanied by progressive changes in tissue microarchitecture.

#### 2.4.3. Duodenum

Figures 15(a)–15(f) illustrate the longitudinal evolution of D-FFOCT dynamic contrast in freshly excised mouse duodenal tissue over 15 h. At baseline, the mucosa exhibits pronounced intracellular dynamic contrast, with clearly distinguishable cellular textures and well-defined glandular and villus-associated structures. Unlike the rapid early reduction in dynamic contrast observed in the brain or the progressive decline in the liver, the duodenum initially retains substantial intracellular contrast, with some regions exhibiting transiently enhanced signals during the first few hours. However, pronounced changes become apparent after approximately 3–5 h, as the intracellular dynamic contrast progressively weakens and the original cellular textures become less distinct. By 8–11 h, many cellular regions exhibit markedly reduced internal contrast, while portions of the surrounding tissue boundaries remain visible. At 15 h, most of the imaged regions display substantially diminished intracellular dynamic signals, with residual contrast primarily associated with larger structural boundaries and localized tissue features. This evolution suggests a delayed but relatively rapid loss of intracellular dynamic contrast, rather than an immediate decline following excision.

Quantitative measurements reveal a corresponding temporal transition. Hmean remains near its baseline value of 55.90 Hz during the first 3 h, followed by a rapid increase to 92.31 Hz at 6.5 h and 112.03 Hz at 15 h (Fig. 15(g)). In contrast, Vmean initially increases from 43.40 to 50.44 a.u. at 1 h and remains above baseline at 3 h, before decreasing sharply to 14.98 a.u. at 6.5 h and 8.80 a.u. at 15 h (Fig. 15(h)), corresponding to an overall reduction of approximately 80%. The most pronounced changes in both parameters occur between approximately 3 and 6.5 h, consistent with the transition from relatively preserved intracellular dynamic contrast to substantially diminished cellular signals observed in the images. Meanwhile, the increasing Hmean indicates a shift toward higher characteristic frequencies among the measured fluctuations, rather than necessarily enhanced cellular activity.

It should be noted that tissue deformation and local displacement affected longitudinal image registration, and the imaging plane at 15 h differed from baseline by approximately 40 µm. Therefore, the observed changes may include contributions from variations in imaging position and tissue geometry. Nevertheless, the consistent temporal trends in Hmean and Vmean across the analyzed regions indicate a pronounced change in the measured dynamic characteristics of the duodenal tissue over the observation period.

**Fig. 15.**
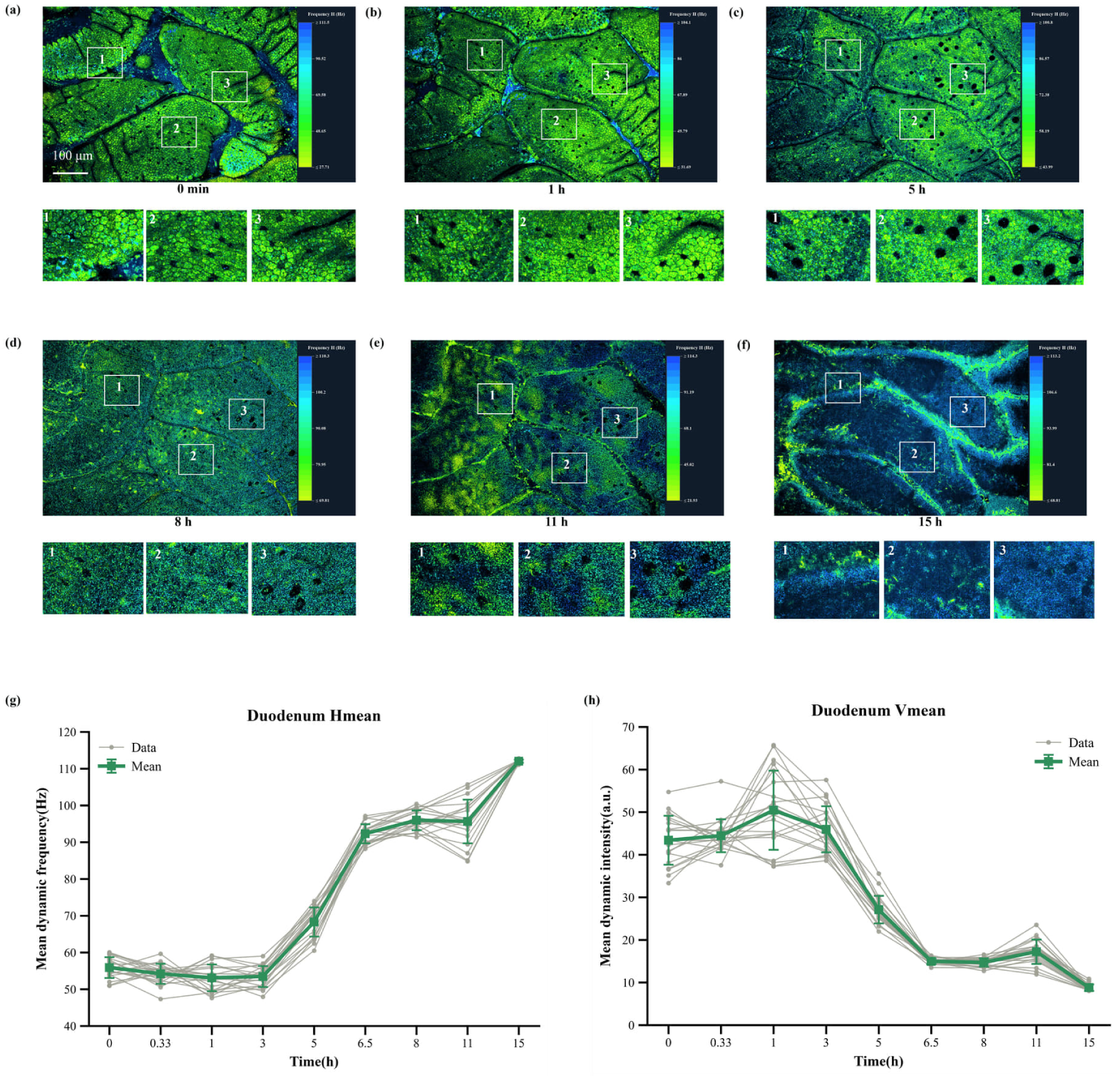
Longitudinal changes in endogenous D-FFOCT dynamics of freshly excised mouse duodenal tissue. (a)–(f) Representative dynamic frequency maps of a corresponding duodenal mucosal region acquired at 0, 1, 5, 8, 11, and 15 h after excision, respectively. Three numbered regions of interest are indicated, with magnified views illustrating temporal changes in local mucosal microarchitecture. (g) Temporal evolution of mean dynamic frequency (Hmean, Hz) over 15 h. (h) Corresponding changes in mean dynamic intensity (Vmean, a.u.). Gray lines represent individual analyzed cellular or local regions (n = 20), and green markers and error bars indicate the mean ± standard deviation. Measurements were obtained at nine time points, displayed at equally spaced positions along the horizontal axis. Tissue deformation and local displacement affected longitudinal registration, and the imaging plane at 15 h was shifted by approximately 40 µm relative to baseline. Scale bar: 100 µm. Scale bar: 100 µm.

#### 2.4.4. Comparison of tissue-dependent temporal dynamics

Comparison of the three tissues reveals distinct temporal patterns of endogenous dynamic contrast following excision. The brain exhibits a rapid early reduction in intracellular dynamic intensity, followed by a prolonged period of relative stability, during which cellular outlines and the overall layered architecture remain partially preserved. In contrast, the liver shows a progressive decline in dynamic intensity throughout the observation period, with intracellular contrast gradually weakening while larger-scale tissue structures remain recognizable. The duodenum displays a different response, characterized by initially preserved or transiently enhanced intracellular dynamic contrast, followed by a rapid reduction after approximately 3–5 h. Despite these differences, all three tissues exhibit an overall increase in Hmean accompanied by a decrease in Vmean, indicating a shift toward higher characteristic fluctuation frequencies and lower dynamic signal amplitudes.

Importantly, the temporal evolution of dynamic contrast does not necessarily coincide with the disappearance of recognizable tissue structures. In the brain, intracellular dynamic signals diminish rapidly while cellular outlines remain visible, whereas the liver and duodenum exhibit more gradual or delayed changes in intracellular contrast. These observations highlight the complementary nature of morphological and dynamic information provided by D-FFOCT. Although the physiological mechanisms underlying these changes remain to be established, the distinct temporal trajectories suggest that different tissues may require organ-specific criteria for longitudinal assessment. Together, these findings demonstrate the potential of D-FFOCT to characterize post-excision tissue changes through combined measurements of dynamic frequency and intensity, providing a basis for future investigations of tissue preservation and viability.

## 3. Materials and methods

### 3.1. Experimental animals and tissue collection

C57BL/6J mice aged 8–12 weeks were used in this study. All animal experiments were approved by the Institutional Animal Care and Use Committee of Tsinghua University (protocol no. 24-XP1). Mice were euthanized by cervical dislocation, followed by immediate tissue dissection. Freshly excised specimens were temporarily maintained in prechilled isotonic phosphate-buffered saline (PBS), and sample mounting and initial image acquisition were completed within 5 min after euthanasia.

Ten organs were examined, including the brain, eyeball, thymus, inguinal lymph node, liver, pancreas, duodenum, kidney, testis, and ovary. Testicular and ovarian specimens were obtained from male and female mice, respectively. These organs were selected to encompass diverse tissue architectures and cellular compositions, including neural, lymphoid, glandular, epithelial, renal, and reproductive tissues. All D-FFOCT measurements were performed on fresh, unfixed, and unstained specimens unless otherwise specified.

### 2.2. Tissue preparation and histological validation

Freshly excised organs were prepared according to their anatomical characteristics to expose the regions of interest. Tissue specimens were immersed in PBS (137 mM NaCl, 2.7 mM KCl, 10 mM Na₂HPO₄, and 2.0 mM KH₂PO₄; pH 7.4) and placed in a custom sample holder with the imaging surface facing upward. A clean coverslip was positioned over the specimen, and the holder was gently closed to flatten the imaging surface while avoiding excessive compression and trapped air bubbles. An immersion medium was subsequently applied above the coverslip for imaging. All measurements were performed at room temperature without active tissue perfusion, medium exchange, or pharmacological intervention during imaging. For longitudinal measurements, specimens remained mounted in the imaging holder throughout the observation period.

Following D-FFOCT imaging, corresponding tissue specimens were processed for histological validation. Samples were fixed in 4% paraformaldehyde, dehydrated through graded ethanol, cleared with xylene, embedded in paraffin, and sectioned for hematoxylin and eosin (H&E) staining. Histological images were acquired using a bright-field microscope. Sections adjacent to the imaged tissue planes were selected for morphological comparison, using anatomical landmarks such as tissue boundaries, vessels, ducts, and characteristic cellular arrangements. Because tissue fixation and processing may introduce shrinkage and deformation, histological and D-FFOCT images were compared qualitatively without pixel-level registration.

### 3.3. Integrated mobile D-FFOCT imaging system

The dynamic full-field optical coherence tomography (D-FFOCT) system (SOX-HPS-L01, SeekOptiX) was based on a low-coherence, full-field Linnik interferometer. Broadband illumination was provided by a light-emitting diode (LED) centered at 660 nm. After collimation and spatial aperture conditioning, the illumination was directed through a Köhler illumination arrangement and divided into reference and sample arms by a 50:50 nonpolarizing beam splitter.

The two interferometer arms incorporated identical water-immersion objectives (UMPLFLN 20×W, Olympus; numerical aperture, NA = 0.5) to provide matched optical configurations. A reflective interface of a yttrium aluminum garnet (YAG) crystal served as the reference reflector. Light returned from the reference and sample arms was recombined to generate full-field interference signals, which were relayed onto a customized complementary metal–oxide– semiconductor (CMOS) camera. The camera operated at 450 frames per second with an array of 1600 × 1100 pixels, without pixel binning.

The system provided a nominal lateral resolution of approximately 0.75 µm and an estimated axial resolution of approximately 3 µm based on coherent length and objective’s NA. The effective field of view was 720 × 495 µm^2^. Specimens were positioned using a motorized three-axis translation stage, enabling automated lateral mosaicking and depth-resolved imaging. Acquisition control, stage movement, image reconstruction, and data processing were coordinated by an industrial computer using central processing unit (CPU) and graphics processing unit (GPU) resources. The optical and computational components were integrated into a mobile cart-based platform measuring approximately 55 × 67 × 100 cm, allowing flexible deployment in tissue preparation and biomedical imaging facilities.

### 3.4. Image acquisition

Before acquisition, the interferometric imaging assembly was adjusted to position the coherence-gated imaging plane within the tissue region of interest. At each field of view, a sequence of N consecutive interferometric frames was recorded for dynamic contrast analysis. The camera operated at 450 frames per second, and 1,024 frames were acquired per measurement in the longitudinal experiments, corresponding to approximately 2.28 s of continuous temporal sampling.

For large-area imaging, an initial sample location was selected as the spatial reference, and the motorized stage sequentially scanned predefined x–y positions with approximately 10% overlap between adjacent fields of view. The acquired images were subsequently combined to generate large-area mosaics. Under the 1,024-frame acquisition condition, approximately 1 cm^2^ of tissue could be scanned within 15 min, including stage movement and acquisition overhead. Shorter acquisition times could be achieved by reducing the number of temporal frames per field of view.

For volumetric imaging, the sample was translated along the axial direction with a step size of 1 µm, and dynamic image sequences were acquired at successive depths. The system supported axial scan ranges exceeding 200 µm, although the effective imaging depth depended on the optical properties of the tissue.

Longitudinal experiments were performed on freshly excised brain, liver, and duodenal specimens using the following acquisition schedules: Brain: 30h; Liver: 20h; Duodenum:15h. Between acquisition time points, the LED illumination was switched off to minimize unnecessary light exposure. The observation periods were selected separately for the three tissue types.

### 3.5. Image processing and volumetric reconstruction

The recorded interferometric sequences were processed using a GPU-assisted computational workflow to obtain dynamic contrast images. For each pixel, the temporal intensity fluctuations were analyzed to extract the dynamic frequency and intensity parameters. The resulting parameter maps were visualized using pseudocolor representations.

For large-area imaging, adjacent fields of view were assembled according to their recorded stage coordinates and overlapping image regions. For volumetric imaging, depth-resolved parameter maps were ordered according to their axial positions to construct three-dimensional image stacks. Volumetric renderings and representative en face images were generated from these stacks.

For longitudinal measurements, corresponding tissue regions were identified at successive acquisition time points. Three representative regions of interest were selected for visualization in each tissue, with magnified views displayed alongside the large-area dynamic images. Quantitative parameters were extracted from 20 selected cellular or local tissue regions per organ. Tissue displacement and changes in the imaging plane were considered when interpreting temporal correspondence, particularly for the duodenal measurements.

### 3.6. Quantification of endogenous dynamic parameters

Two quantitative parameters were extracted from temporal interferometric intensity fluctuations: mean dynamic frequency (Hmean) and mean dynamic intensity (Vmean). For each pixel, the recorded temporal intensity sequence I(t) was normalized, and its one-sided power spectral density P(f) was calculated. Spectral analysis was performed over the frequency range of 0.5–225 Hz, with the upper limit corresponding to the Nyquist frequency of the 450-Hz acquisition. The mean dynamic frequency (Hmean) was defined as the power-weighted average frequency within the analyzed spectral range: Hmean = Σ[f·P(f)] / ΣP(f)

The mean dynamic intensity (Vmean) was calculated by integrating the power spectral density over the same frequency range: Vmean = Σ[P(f)·Δf] where fmin = 0.5 Hz, fmax = 225 Hz, and Δf denotes the frequency-bin width. Hmean characterizes the temporal frequency distribution of interferometric fluctuations and is expressed in hertz (Hz). Vmean quantifies the total spectral power associated with temporal intensity fluctuations and is reported in arbitrary units (a.u.).

Both parameters were calculated at the pixel level to generate spatially resolved dynamic maps. For cross-organ comparisons, Hmean and Vmean were averaged within individual imaging fields, with each field treated as one measurement location. For longitudinal experiments, the parameters were extracted from selected cellular or local tissue regions at successive acquisition time points. Together, Hmean and Vmean provide complementary quantitative descriptions of endogenous interferometric fluctuations, characterizing their frequency distribution and magnitude-related spectral power, respectively.

### 3.7. Statistical analysis

Quantitative results are presented as mean ± standard deviation (SD), unless otherwise specified. Box plots display the median and interquartile range, with individual measurement locations represented by overlaid data points. For cross-organ comparisons, Hmean and Vmean were analyzed across 120 imaging locations from nine organs: brain, eyeball, thymus, inguinal lymph node, liver, pancreas, duodenum, kidney, and testis. Each organ contributed 9–17 imaging locations. The ovary was included in the structural atlas but not in the cross-organ quantitative comparison. Differences among organs were evaluated using one-way analysis of variance (ANOVA), followed by Tukey’s multiple-comparison test for pairwise comparisons. A significance threshold of p < 0.05 was used. For longitudinal measurements, temporal Hmean and Vmean values from 20 selected cellular or local tissue regions per organ were summarized at each acquisition time point. Individual trajectories and group means with standard deviations were displayed to characterize temporal changes. Imaging locations and tracked regions were treated as measurement units. Where multiple measurements originated from the same animal or tissue specimen, they were not interpreted as independent biological replicates.

## 4. Discussion

In this study, we established a label-free volumetric dynamic atlas of ten fresh mouse organs using an integrated D-FFOCT platform. By combining large-area imaging, depth-resolved reconstruction, and quantitative dynamic analysis, this work provides a systematic reference for tissue characterization across diverse organs. Large-area mosaics reveal characteristic anatomical structures, while volumetric imaging provides additional information about local three-dimensional microarchitecture. The level of morphological detail varies among organs, reflecting differences in cellular organization and optical properties. Compared with conventional histology and tissue-clearing methods, D-FFOCT enables direct examination of fresh tissues without fixation, staining, or physical sectioning, preserving specimens for subsequent analyses. Although it cannot replace the molecular specificity of established atlas technologies, it provides complementary structural and dynamic information under native tissue conditions.

Beyond morphological characterization, quantitative analysis revealed distinct endogenous dynamic signatures across nine organs. The mean dynamic frequency (Hmean) and mean dynamic intensity (Vmean) exhibited different organ-dependent distributions. Notably, the pancreas showed relatively high fluctuation frequencies but low dynamic intensity, whereas the liver exhibited the opposite behavior. These findings suggest that Hmean and Vmean capture complementary aspects of tissue dynamics rather than representing equivalent measurements. Such differences may reflect variations in intracellular motion, cellular organization, and optical scattering properties. However, the underlying biological mechanisms remain to be established, and these parameters should not be interpreted as direct measurements of metabolic activity. Our results provide a comparative reference for understanding tissue-dependent dynamic contrast and suggest that combining frequency- and intensity-based measurements may improve quantitative tissue characterization.

Longitudinal imaging further demonstrated distinct temporal changes in freshly excised tissues. Brain tissue exhibited a rapid early reduction in dynamic intensity followed by relative stabilization, whereas liver tissue showed a more gradual and sustained decline. Duodenal tissue initially maintained substantial dynamic contrast before undergoing a pronounced reduction after approximately 3–5 h. Despite these differences, all three tissues exhibited increasing Hmean and decreasing Vmean over time. Importantly, the loss of intracellular dynamic contrast did not necessarily coincide with the disappearance of recognizable tissue structures, suggesting that dynamic imaging may reveal tissue changes that are less apparent in morphological images alone. Nevertheless, the increasing Hmean should not be interpreted as enhanced cellular activity, and further experiments combining independent viability and metabolic measurements are required to clarify the physiological significance of these observations.

Several limitations remain. The effective imaging depth is restricted by tissue scattering, limiting volumetric characterization to relatively superficial regions. Histological comparisons are affected by deformation during tissue processing, while quantitative measurements from selected imaging locations may not fully represent biological variability across animals. Tissue displacement and changes in imaging depth may also influence longitudinal measurements. Future studies involving larger animal cohorts, improved image registration, and independent functional validation will help establish more reliable quantitative references. Importantly, the publicly available raw interferometric datasets from all ten organs provide a resource for reproducible analysis, algorithm development, and further methodological comparisons. By integrating large-area morphology, volumetric microarchitecture, and endogenous dynamic measurements, this atlas provides a foundation for extending D-FFOCT toward systematic tissue characterization under different physiological and pathological conditions.

## 5. Conclusion

In this study, we established a label-free volumetric dynamic atlas of ten fresh mouse organs using an integrated mobile D-FFOCT imaging platform. By combining large-area imaging, depth-resolved reconstruction, and quantitative analysis of endogenous dynamics, we systematically characterized organ-specific tissue microarchitecture and dynamic signatures. Cross-organ comparisons revealed distinct distributions of dynamic frequency and intensity, while longitudinal measurements demonstrated tissue-dependent changes in endogenous dynamics following excision. Importantly, dynamic contrast changes could occur while recognizable tissue structures remained preserved, highlighting the complementary information provided by morphological and dynamic imaging. Furthermore, the publicly released raw interferometric datasets provide an accessible resource for reproducible analysis and future methodological development. Together, this work establishes a comprehensive optical reference of fresh mouse organs and provides a foundation for extending D-FFOCT to systematic tissue characterization and investigations of physiological and pathological changes.

## Supporting information

Visualization 1. Large-area D-FFOCT imaging of mouse brain (Fig. 2a)

Visualization 2. 3D D-FFOCT imaging of mouse hippocampus (Fig. 2d)

Visualization 3. 3D D-FFOCT imaging of mouse cornea (Fig. 3d)

Visualization 4. 3D D-FFOCT imaging of mouse liver (Fig. 6d)

Visualization 5. Mouse dentate gyrus time-lapse over 30 h (Fig. 13a-f)

Visualization 6. Mouse liver time-lapse over 20 h (Fig. 14a-f)

## 7. Acknowledgment

This work was supported by the National Key Research and Development Program of China (2024YFC3405300, 2025YFC2422701).

## 9. Data availability

The raw imaging data and high-resolution, large-area images of each organ are publicly available at https://github.com/forest1997/mouse-organs-dffoct-raw-data. Instructions for downloading, reconstructing and verifying the data are provided in the repository. These datasets include the original time-resolved interferometric image sequences used for morphological visualization and quantitative dynamic analysis. See Visualizations 1–6 for supporting content.

